# Rice JASMONIC ACID OXIDASES (OsJAO) control resting jasmonate metabolism to promote development and repress basal immune responses

**DOI:** 10.1101/2024.07.24.604933

**Authors:** Simon Ndecky, Ludivine Malherbe, Claire Villette, Véronique Chalvon, Isabelle Meusnier, Dennisse Beltran-Valencia, Nicolas Baumberger, Michael Riemann, Thomas Kroj, Antony Champion, Thierry Heitz

## Abstract

Recent research has established that catabolic conversions within the jasmonate pathway have significant consequences on hormone signaling output. In dicotyledonous plants, the jasmonic acid oxidase (JAO) catabolic route is endowed with a regulatory function by diverting jasmonic acid (JA) towards hydroxylation, at the expense of its conjugation into the bioactive jasmonoyl-isoleucine (JA-Ile) hormone. Here we functionally characterized the JAO pathway in rice (*Oryza sativa*) and demonstrate its prevalent function in promoting growth and attenuating JA responses in vegetative tissues. The rice genome contains four JAO-related homologs of which three generated hydroxy-JA *in vitro* and reverted the high defense phenotype when expressed in the Arabidopsis *jao2-2* mutant. By generating and analyzing a series of single to quadruple rice *jao* mutants, we show the incremental effect of gradual JAO depletion on JA metabolism, basal defense levels, growth inhibition, fitness and global metabolic reprogramming. JAO-deficient lines were significantly growth-retarded at the juvenile stage, while recovering a near wild-type vegetative development after three months, where they exhibited a enhanced resistance to virulent and avirulent strains of *Magnaporthe oryzae*, the causal agent of fungal blast disease. Our findings identify the JAO pathway as an integral component of rice JA homeostasis and an important determinant of the growth-defense tradeoff. They demonstrate its conserved regulatory function in monocots and open possibilities for modulating selectively basal JA responses in a major cereal crop. Natural variation in JAO activity could also be explored as a mechanism underlying varying levels of JA signaling output in rice.

## Introduction

Jasmonates (JAs) are fatty acid-derived compounds with regulatory functions that impact the entire plant life cycle, from germination to vegetative growth and reproductive success. JAs have critical functions in the interaction of plants with their biotic environment including beneficial and pathogenic organisms, and largely orchestrate a conserved antagonism between growth and defense (Wasternack and Hause, 2013; Campos et al., 2014; Guo et al., 2018). JAs are best characterized as positive regulators of transcriptional responses to herbivores or to microbial pathogens, but recent efforts have uncovered additional functions in the adaptation to numerous abiotic stresses including drought, salt, extreme temperatures or nutrient stress (Kazan, 2015; Ali and Baek, 2020; Raza et al., 2020).

Synthesis of jasmonic acid (JA), the name-giving compound and hormone precursor, is initiated in plastids upon release of linolenic acid from membrane lipids (Kimberlin et al., 2022), before its oxygenation to an allene oxide intermediate that can be cyclized into the first jasmonate in the pathway, 12-oxo-phytodienoic acid (OPDA). In the major biosynthetic route, a fraction of OPDA is converted to JA in the peroxisomes that will be transferred in the cytosol (Wasternack and Hause, 2013; Heitz, 2020). Even though exogenous JA triggers extensive transcriptional reprogramming and physiological responses, this precursor acquires its activity only endogenously through JASMONATE RESISTANT 1 (JAR1)-catalyzed conjugation into jasmonoyl-isoleucine (JA-Ile) (Staswick and Tiryaki, 2004), which is one among numerous possible metabolic fates (Miersch et al., 2008; Heitz et al., 2019).

Under low hormone levels, JA-responsive transcription factors (TFs) are kept inactive by a family of JASMONATE ZIM-DOMAIN (JAZ) repressor proteins and associated co-repressors (Pauwels and Goossens, 2011). When developmental- or stress-induced JA-Ile formation occurs, the conjugate acts as a ligand that promotes the assembly of receptor complexes whose core components are the F-box protein CORONATINE INSENSITIVE 1 (COI1) and various JAZ protein(s). COI1-JAZ complexes then engage into E3 ubiquitin ligases which will proteolytically degrade ubiquitinated JAZ, resulting in the de-repression of target transcription (Chini et al., 2007; Thines et al., 2007).

In addition to complex protein interactions in signaling processes, insights during the last decade have established that hormone turnover and other metabolic modifications within the JA biochemical pathway significantly contribute to fine tuning JA-Ile response output. Knowledge initially gained in Arabidopsis has defined a complex metabolic grid (Fig. 1A) where JA-Ile catabolism is central to generate peculiar JA blend signatures in specific organs or physiological situations (Widemann et al., 2016; Heitz et al., 2019). After its formation and perception, JA-Ile is rapidly and simultaneously turned over by oxidative and deconjugation pathways. Oxidation is catalyzed by three JA-regulated cytochrome P450 of the CYP94 subfamily that generate the less active 12OH-JA-Ile and inactive 12COOH-JA-Ile derivatives (Kitaoka et al., 2011; Koo et al., 2011; Heitz et al., 2012; Jimenez-Aleman et al., 2019; Poudel et al., 2019). JA-Ile as well as its first catabolite 12OH-JA-Ile are also substrate of the amido-hydrolases (AH) IAR3 and ILL6 to generate free JA and 12OH-JA respectively (Widemann et al., 2013; Zhang et al., 2016). The genetic disruption of these metabolic pathways strongly modifies JA-Ile homeostasis with increased steady-state levels and enhanced half-life while their ectopic overexpression depletes the hormone and attenuates downstream responses (Koo et al., 2011; Heitz et al., 2012; Widemann et al., 2013; Zhang et al., 2016; Marquis et al., 2020). Unexpectedly, in Arabidopsis, impaired JA-Ile catabolism does not systematically result in enhanced defense signaling, as such a ‘super-defense’ scenario seems to occur only in some pathway- and stress type-specific combinations, in which repressor feed-back mechanisms are not themselves overstimulated (Marquis et al., 2020). In contrast, a more linear output was described in wild tobacco where silencing of a set of CYP94 enzymes efficiently boosted defense and plant resistance against a generalist herbivore (Luo et al., 2016).

**Figure 1.**
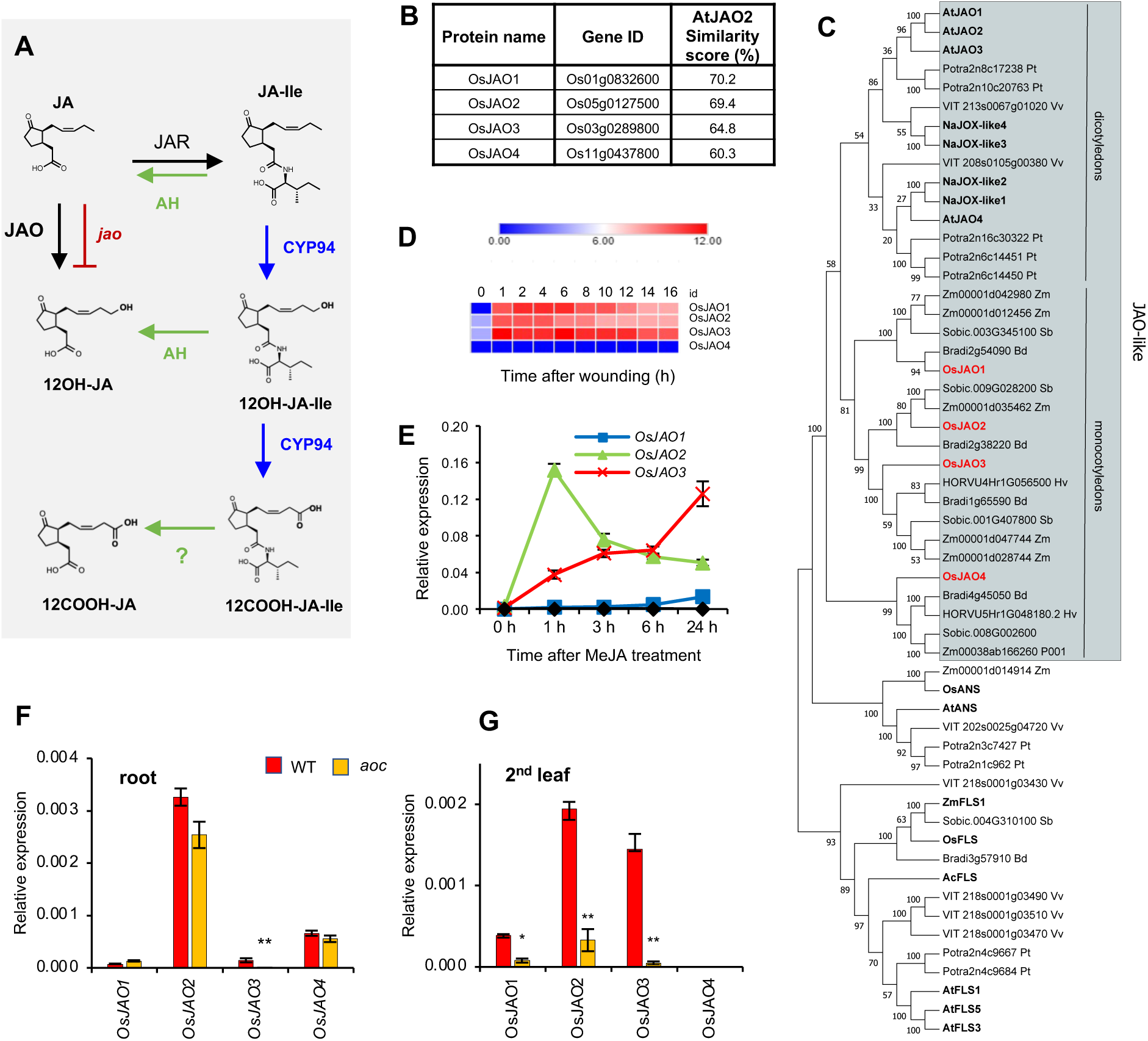
JA/JA-Ile catabolic pathways and identification of candidate *JAO* genes in rice. (a) Simplified metabolic grid of JA/JA-Ile catabolic pathways as initially elucidated in Arabidopsis. AH: amido-hydrolase; CYP94: cytochrome P450 family 94; JAO: jasmonic acid oxidase; JAR: jasmonate- resistant. Question mark depicts an uncharacterized step. (b) Nomenclature, locus ID (RAPdb) and similarity score of the four predicted *OsJAO* proteins to Arabidopsis JAO2 protein. (c) Phylogenic tree of AtJAO2-related proteins in dicot and monocot plants. Characterized proteins in *A. thaliana* (At) and *N. attenuata* (Na) were displayed along with sequences sharing > 35 % sequence identity with AtJAO2 from representative species in dicots (*Populus tremula* Potra, *Vitis vinifera* Vv) and monocots (*Brachypodium distachyon* Bd, *Hordeum vulgare* Hv, *O. sativa* Os, *Sorghum bicolor* Sobic, *Zea mays* Zm). Bootstrap values after 100 iterations are shown on nodes. Protein alignment used for constructing tree is shown in Supplemental Figure S1. (d) Kinetic profile of *OsJAO* gene expression in a time series of leaf wounding as obtained by RNAseq. (e) Kinetic profile of *OsJAO* gene expression in leaves of 10-day old seedlings after exposure to volatile MeJA. (f) and (g) Basal expression levels of *OsJAO* genes in leaves (f) or roots (g) of wild-type (WT) or JA-deficient *aoc* plants. For (e), (f) and (g), expression was determined by RT-qPCR and was normalized with signal from housekeeping genes *UBQ5* and *UBQ10*. (d) to (g) Data display means of three biological replicates with SEM. (f) and (g) Asterisks indicate a significant difference between genotypes as determined by t-test (\**P*<0.05; \*\**P*<0.01).

More upstream in the biochemical pathway, diverse JA modifications are known as hydroxylation, sulfation, glucosylation, decarboxylation or conjugation (Wasternack and Hause, 2013); however, their mode of formation or physiological relevance beyond storage was unknown until recently. For example, formation of sulfated or glycosylated JA derivatives is common in plants (Miersch et al., 2008) and requires prior hydroxylation. In addition to deconjugation of 12OH-JA-Ile by the AH pathway (Widemann et al., 2013), a direct JA hydroxylation pathway was deciphered in Arabidopsis in the form of four JASMONIC ACID OXIDASES (JAO/JOX, Fig. 1A and 1C) that belong to a JA-regulated subclade of 2-oxoglutarate-dependent oxygenases. Unexpectedly, the specific suppression of the AtJAO2 isoform is sufficient to redirect metabolic flux towards JA-Ile formation and enhance constitutive JA-Ile-directed responses and antifungal resistance (Smirnova et al., 2017). Multiple *jao* mutations further strengthen insect attack tolerance and drought survival phenotypes (Caarls et al., 2017; Marquis et al., 2022). These findings highlight the peculiar and potent regulatory function of the JAO pathway by defining a metabolic sink impacting JA-Ile formation and action. Consequently, varying the extent of JA hydroxylation activity may be a means plants use to modulate JA-Ile signaling and this feature deserves to be explored in species of agronomic interest.

Rice (*Oryza sativa*) is a crop species of uppermost importance as the major staple food crop for more than half of the world’s population and has been developed as a model in cereal research with ever increasing resources and technologies. Given its wide genetic diversity and cultivation in diverse habitats, rice is exposed to a tremendous range of biotic and abiotic stresses threatening its productivity. In this context, the rice JA hormonal pathway has been extensively investigated, revealing many conserved and some specific features (Dhakarey et al., 2016; Nguyen et al., 2019). For example, some prominent functions of JA in rice are in spikelet development and anther dehiscence (Cai et al., 2014; Xiao et al., 2014), antifungal (Mei et al., 2006; Riemann et al., 2013) or anti-herbivore (Xu et al., 2021; Zeng et al., 2021) defense. In rice, JA-Ile is perceived by functionally diversified COI receptors that have specialized in regulating partially distinct responses. Notably, the divergent COI2 protein harbors important functions in mediating root growth regulation, male fertility, senescence and antimicrobial defense (Inagaki et al., 2023; Nguyen et al., 2023; Wang et al., 2023). Orthologues of most JA biosynthetic and signaling genes have been identified in rice and were functionally characterized in some cases (Nguyen et al., 2019). In contrast, knowledge on jasmonate catabolism and its physiological role is scarce in this species. Rice homologs of CYP94 and AH enzymes and their enzymatic products have been characterized in response to leaf wounding and salt stress (Hazman et al., 2019). Among a series of stress tolerance genes, the most striking gain in salt tolerance was by overexpressing a JA-Ile deactivating enzyme (Kurotani et al., 2015), consistent with the finding that JA signaling is globally detrimental to salt tolerance (Ndecky et al., 2023). Moreover, natural variation in CYP94- catalyzed JA-Ile catabolism was reported in rice and haplotypes with weaker expression of CYP94C2 where characteristic of accessions better adapted to a temperate climate (Mao et al., 2019). These data provide a link between catabolic plasticity in the JA pathway and peculiar phenotypic characteristics.

To further explore metabolic regulation of JA signaling and its potential to modulate agronomic traits, we set out to investigate the JAO pathway in rice. Out of four *AtJAO2*-related rice genes, three were found to be JA-regulated and to encode JAO activity. By developing and analyzing a collection of single to quadruple rice os*jao* mutant lines, we show that JAO activity depletion incrementally shifts jasmonate metabolism towards enhanced JA-Ile formation and catabolism. Phenotypically, os*jao* mutants display repressed juvenile growth, elevated basal expression of JA marker genes, substantial metabolic reprogramming, and increased resistance to the blast fungal disease caused by *Magnaporthe oryzae*. Our study uncovers the need to tightly control basal JA homeostasis to repress costly JA responses and optimize growth in a cereal species.

## Results

### Rice expresses four *AtJAO2*-related genes

By using AtJAO2 protein as a query, four related sequences were retrieved from the predicted proteome of the rice reference accession Nipponbare (IRGSP-1.0 annotation). According to decreasing similarity with AtJAO2, they were tentatively named OsJAO1 (Os01g0832600), OsJAO2 (Os05g0127500), OsJAO3 (Os03g0289800) and OsJAO4 (Os11g0437800) (Fig. 1B). When constructing a phylogenic tree of sequences originating from representative species from angiosperm lineages, OsJAO sequences formed a monophyletic clade with other monocot sequences in the DOX46 subclade of 2-oxoglutarate/Fe (II)-dependent dioxygenases (2-ODDs) defined by (Kawai et al., 2014), adjacent to a group of dicot proteins including JAOs characterized functionally from Arabidopsis (Caarls et al., 2017; Smirnova et al., 2017) and *Nicotiana attenuata* (Tang et al., 2020) (Fig. 1C; Supplementary Fig. S1). The JAO-related subclade of ODDs clustered close to another group formed of flavonoid biosynthetic genes from both monocots and dicot species.

As characterized dicot *JAO* genes are notoriously co-regulated with JA pathway genes (Caarls et al., 2017; Smirnova et al., 2017), we tested the response of the *OsJAO* genes in leaves to either mechanical wounding or MeJA treatment. *OsJAO4* transcripts were not detected in leaves in any tested condition, whereas *OsJAO1*, *2* and *3* expression was rapidly induced after wounding, reaching a plateau by 1 h throughout at least 16 h (Fig. 1D). In response to MeJA exposure, contrasted profiles were recorded: *OsJAO2* transcripts peaked early by 1 h before declining, *OsJAO3* increased steadily until 24 h and *OsJAO1* showed a weak and late increase (Fig. 1E). Because basal expression level rather than inducibility was demonstrated to determine biological function in Arabidopsis (Smirnova et al., 2017), we examined precisely the relative transcript levels of *OsJAO* in unstressed rice tissues in WT and JA-deficient *aoc* mutant (Nguyen et al., 2020). *OsJAO2* expressed the highest in roots, followed by *OsJAO4*, whereas *OsJAO3* and *OsJAO1* only showed marginal signal (Fig. 1F). Of note, all genes except *OsJAO3* were equally expressed in both genotypes. In leaf material, *OsJAO2* and *OsJAO3* dominated *OsJAO1* expression, and all 3 genes lost most basal expression in *aoc* (Fig. 1G). These expression data were consistent with a possible role of *OsJAO* genes in JA metabolism.

### Rice *OsJAOs* encode active jasmonic acid oxidases

Alignment of the predicted OsJAO protein sequences with those of JA oxidases from Arabidopsis and *N. attenuata* revealed conservation of almost all residues essential for the binding of the substrates JA and 2-oxoglutarate in structure/function analysis of AtJOX2/JAO2 (Zhang et al., 2021). The only exception was that at position 308 in OsJAO4, the conserved serine (Ser) was substituted by threonine (Thr) (Supplementary Fig. S2). These features suggest that *OsJAO* genes encode active JA oxidases. To test this hypothesis, recombinant OsJAO proteins were expressed in bacteria and clarified lysates or purified proteins were assayed for jasmonic acid oxidase activity with 2-oxoglutarate (2OG) as a co-substrate. Reaction products were analyzed by liquid chromatography coupled to mass spectrometry (LC-MS/MS). All OsJAO proteins, except OsJAO4, produced in the presence of JA and 2OG, a compound presenting the characteristics (retention time and mass spectrum) of an authentic 12OH-JA standard (Fig. 2A). The controls, no-JAO protein or absence of 2OG co-substrate, yielded no oxidized product (Supplementary Fig. S3). An OsJAO4 variant, where the Thr at position 308 was substituted by Ser (OsJAO4p308T>S), also exhibited no JAO activity *in vitro*, indicating that lack of activity of OsJAO4 is not, or not only due to this substitution (Fig. 2A).

**Figure 2.**
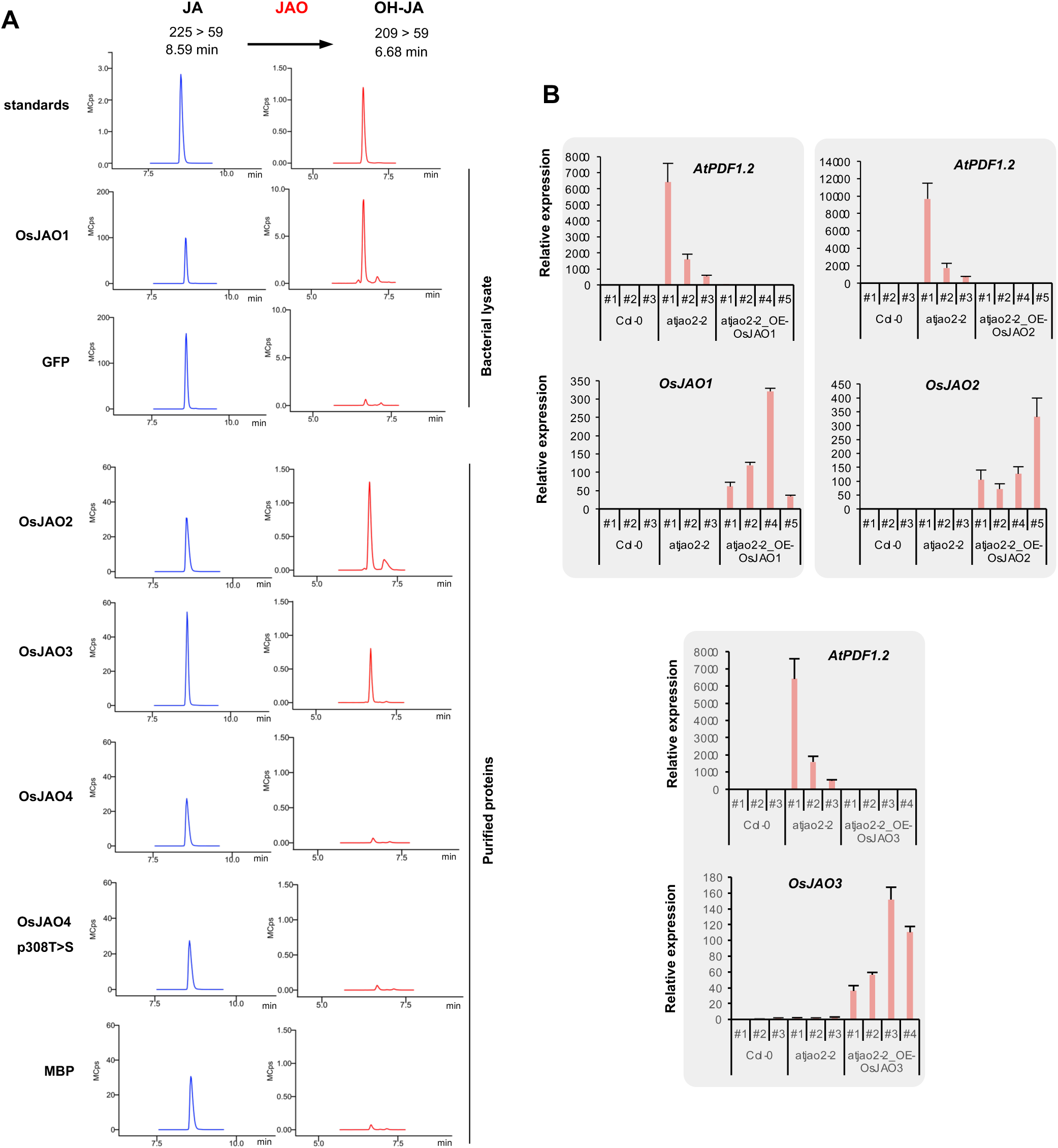
Functional analysis of *OsJAO* genes and proteins. (a) Catalytic activity of recombinant OsJAO proteins on JA. LC-MS/MS chromatograms of residual JA or produced OH-JA are shown in incubation reactions of JA with bacterial lysate (expressing OsJAO1 or GFP) or affinity-purified proteins (MBP, OsJAO2, OsJAO3 or OsJAO4). Incubations were performed in the presence of the co-substrate 2-oxoglutarate. Control reactions in the absence of the co-substrate are shown in Supp Fig. S3. The oxidation product matched the retention time and the mass transition of authentic 12OH-JA standard. (b) *In planta* complementation assay in Arabidopsis *jao2-2* mutant background expressing *OsJAO* (OE). Three control (Col0 WT and *jao2-2*) and four independent T1 transformed plants were analyzed at 3 weeks for impact of *OsJAO* transgene expression on *PDF1.2* marker transcript levels in leaves as determined by RT-qPCR. Expression was normalized with signal from *EXP* and *TIP41* housekeeping genes. Histograms show mean ± SD of 3 measurements of each plant.

Arabidopsis lines impaired in *AtJAO2* expression display an upregulation of JA-dependent defense gene expression in leaves due to a redirection of metabolic flux towards JA-Ile formation and signaling (Smirnova et al., 2017). Taking advantage of this molecular phenotype, we expressed each *OsJAO* gene in the *Atjao2-2* background and assayed *AtPDF1.2* defense marker expression as an *in vivo* readout of JAO function. In Arabidopsis lines expressing OsJAO1, 2 or 3, *AtPDF1.2* expression was reverted back to low, WT levels (Fig. 2B), while for OsJAO4-expressing lines, no consistent data were obtained. These results indicate that OsJAO1, 2 and 3 modify JA homeostasis and signaling in the Arabidopsis *jao2-2* mutant, suggesting functional conservation between the Arabidopsis and rice proteins.

### Generation of a series of single *jao2* and multiple *jao* rice mutants

To investigate the *in vivo* functions of OsJAO enzymes, we generated loss-of-function rice mutant lines using the CRISPR-Cas9 gene editing technology. Based on expression data that initially identified *OsJAO1* and *OsJAO2* as the most expressed genes in leaves, we selected them for generating single mutants using two sgRNAs for each gene (Supplementary Fig. S4A). In addition, we prepared a multiplex construct containing one specific gRNA for each *OsJAO* gene interspersed with tRNA^Gly^ sequences (Xie et al., 2015). Supplementary Fig. S4B shows the positions of each sgRNA in the respective gene structures. For an unresolved reason, all plants regenerated after transformation with the *OsJAO1* construct exhibited a WT *OsJAO1* sequence, and consequently, no single *Osjao1* ko lines were obtained. Homozygous T1 plants were recovered with one (*jao2* #27) or two (*jao2* #55) nucleotide deletions, or one nucleotide insertion in the *JAO2* coding sequence (Supplementary Fig. S4A). From the multiplex construct, homozygous lines with characterized mutations in multiple *JAO* genes were generated and were propagated to T3, Cas9-free populations. These various mutations introduce premature stop codons and the synthesis of truncated, inactive proteins lacking essential residues for binding of Fe^2+^ or the 2OG co-substrate (Supplementary Fig. S4B). In some instances, deletions of 3 nucleotides that change two non-conserved amino acid residue were observed, as for example, in *OsJAO3:p.61PD>H*. Ectopic expression of this mutated protein in the Arabidopsis *jao2-2* mutant reverted At*PDF1.2* expression to WT levels, suggesting it has retained JAO activity (Supplementary Fig. S6). Similarly, an in-frame deletion of 3 nucleotides in *OsJAO4* is unlikely to alter activity, if any. The lines *jao1.2 #67* and *jao1.2.3 #11* were thus considered as double and triple mutants respectively (Supplementary Fig. S4A). Similarly, given these features, lines *jao1.2.4 #137* and *jao1.2.3 #13* were considered as double and triple mutant lines, respectively. Finally, two quadruple mutant lines with frameshift mutations in all four genes were obtained, and named *jao1.2.3.4 #249* and *jao1.2.3.4 #170*. Informative mutants were further analyzed for their macroscopic or molecular phenotypes.

### Rice *jao* mutants exhibit altered jasmonate homeostasis

To evaluate the impact of *jao* mutations on basal JA homeostasis, we quantified six JAs in shoots of 9-d old seedlings of mutant plants grown in the absence of any applied stress. In WT Kitaake plants, basal JA content was around 2 pmol g FW^-1^, JA-Ile was below detection limit, but oxidized derivatives OH-JA, 12OH-JA-Ile, 12COOH-JA-Ile and 12COOH-JA were readily detected in the range of 15-40 pmol g FW^-1^ (Fig. 3). Steady-state levels of JA, the JAO enzyme substrate, were slightly increased in single and double mutants and more consistently in higher order (mostly in triple) mutants. Conversely, abundance of OH-JA, the direct JAO oxidation product, was not affected in single and double lines, but was strongly reduced in triple and quadruple mutant lines. The active hormone JA-Ile was detected in all mutants, with a tendency for more abundance in higher order mutant lines, and these latter accumulated less of its direct catabolite 12OH-JA-Ile, which is a highly turned-over intermediate in the pathway (Heitz et al., 2012). The carboxylated derivatives, free or Ile-conjugated, which are the most downstream and stable JA catabolites known, over accumulated in triple and quadruple mutants. These quantitative alterations in mutant lines are consistent with the position of JAO in the JA metabolic grid (Fig. 1A) and a role of OsJAO enzymes in basal JA homeostasis. They show that cumulative genetic JAO suppression depletes OH-JA formation and promotes JA-Ile accumulation and catabolism in young rice leaves.

**Figure 3.**
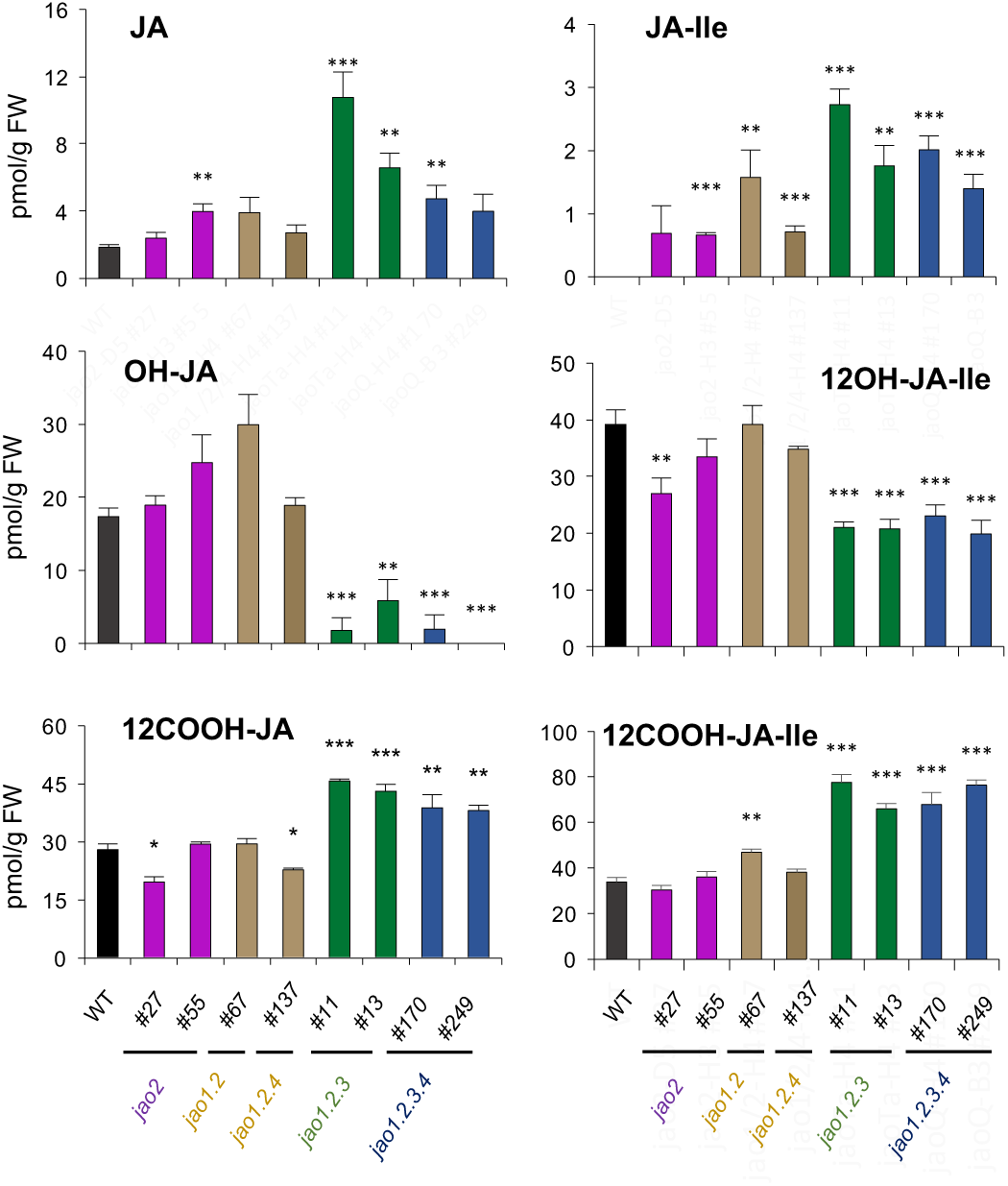
Impact of *jao* mutations on JAs profiles. JAs profiles were analyzed in shoots of WT and a series of *jao* mutants collected from unstressed 9 day-old T3 rice seedlings. Plants were grown on phytoagar in Magenta boxes. JAs were extracted and analyzed by LC-MS/MS. Histograms represent means of 5 biological replicates with SEM. Asterisks represent statistical significance for each mutant line compared with WT (t-test, \**P*<0.05; \*\**P*<0.01; \*\*\**P*<0.001).

### Rice *jao* mutants show developmental phenotypes reminiscent of elevated jasmonate signaling

Jasmonate signaling is well-known to affect early seedling development in rice (Riemann and Takano, 2008; Svyatyna et al., 2014). For example, under dark conditions, JA-deficient rice (*aoc*) develops an exaggerated long mesocotyl relative to WT, illustrating the repressive action of JA signaling on WT mesocotyl elongation (Ndecky et al., 2023) (Fig. 4A). In this assay, the *jao1.2.3.4 #249* displayed significantly shorter mesocotyls than WT, suggesting that loss of JAO activity may result in increased JA signaling and additional repression of mesocotyl development in the dark.

**Figure 4.**
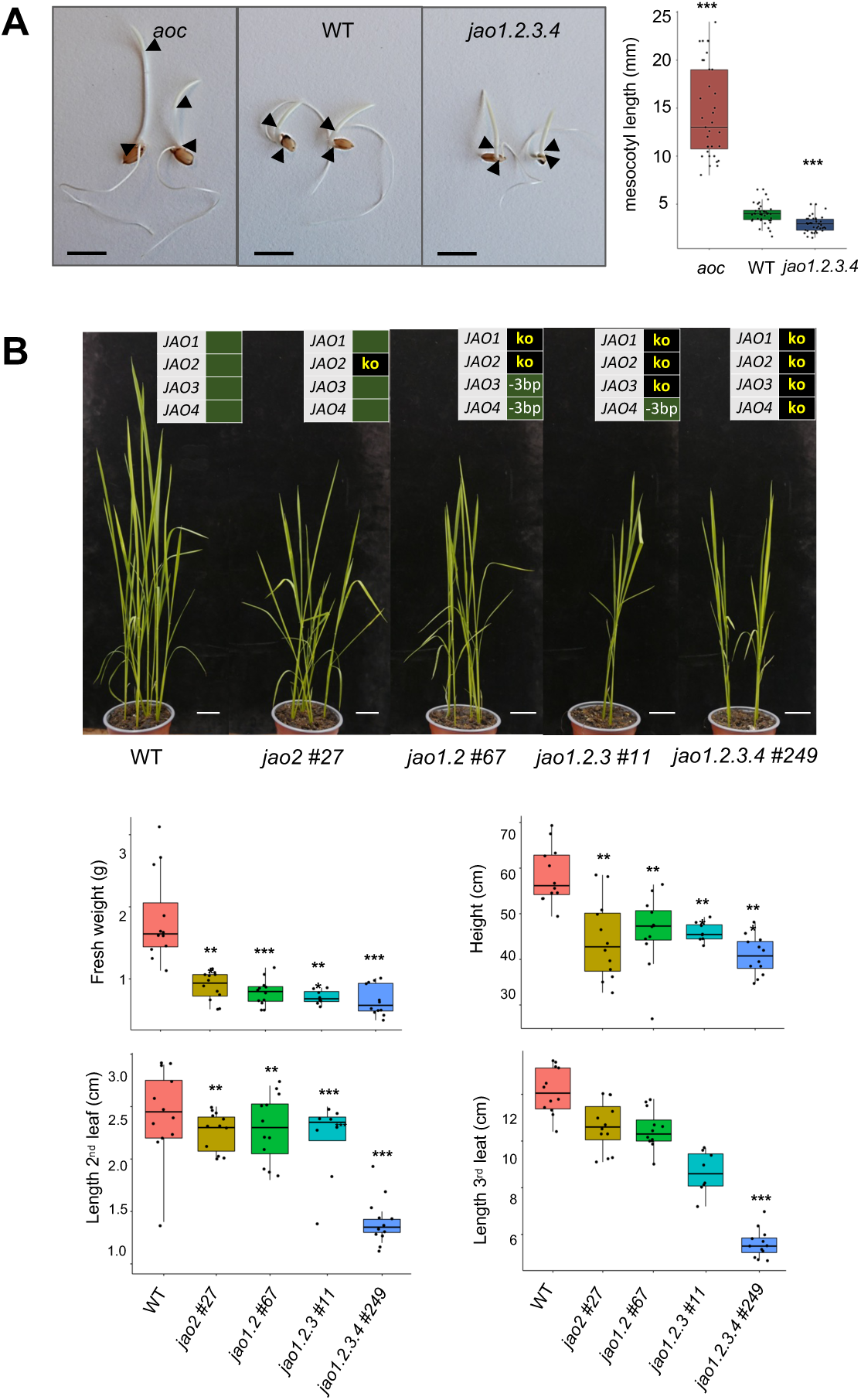
Analysis of growth parameters of rice *jao* mutant lines. (a) Elongation of mesocotyl under dark conditions in WT, JA-deficient (*aoc*) and *jao1.2.3.4 #137* lines. Seeds were germinated in darkness for 4 days in Petri dishes and two representative seedlings are shown in left panel. *aoc/aoc* plants were phenotypically identified from an *AOC/aoc* segregating population by their exaggerated mesocotyl elongation. Black arrowheads indicate limits of the mesocotyl. Scale bar = 1 cm. Right panel represents measurement of mesocotyl length of 35 seedlings for each genotype. Statistical significance relative to WT was assessed by t-test (***P < 0.001). (b) Shoot growth phenotypes of T3 plants of single *jao2*, double *jao1.2*, triple *jao1.2.3* and quadruple *jao1.2.3.4* rice mutants at four weeks after sowing (n=10-12 per genotype). Indicated genotypes were assessed quantitatively for shoot height, shoot fresh weight, 2^nd^ and 3^rd^ leaf length and displayed in lower panels. Scale bar: 2 cm. Asterisks indicate statistical difference with WT (t-test, \*\**P*<0.01; \*\*\**P*<0.001).

Because enhanced basal JA signaling resulted in growth retardation in Arabidopsis *jao* multiple mutants (Caarls et al., 2017; Marquis et al., 2022), we next analyzed rice *jao* mutants for growth characteristics. An initial examination of vegetative development (4 weeks) with T2 generation *jao2* and *jao1.2.3.4* lines under optimal growth conditions indicated a reduction in plant height, shoot biomass, and leaf length, particularly in the quadruple *jao1.2.3.4 #249* mutant (Sup Fig. S7). In a second independent experiment including two additional genotypes, all in T3 generation, a pronounced reduction in plant height and shoot fresh weight was evidenced for single *jao2,* double *jao1.2,* triple *jao1.2.3* and quadruple *jao1.2.3.4* lines (Fig. 4B). Length of leaf 2 was particularly affected, with an incremental negative impact when mutations were cumulated. These observations indicate that JAO deficiency may activate JA signaling in elongating tissues leading to reduced vegetative growth.

Two independent single *jao2* and two *jao1.2.3.4* lines were next surveyed at a later developmental stage. By 3 months, the vegetative growth differences were less prominent and comparable between mutant genotypes (Supplementary Fig. S8), suggesting a declining impact of modified JA metabolism throughout rice vegetative development. At the reproductive stage, we observed in all analyzed mutants, a decrease in fertility that was however only significant in one quadruple *jao* mutant line, and a reduction of seed mass by about 20% (Supplementary Fig. S8).

### JAO deficiency results in elevated basal defense gene expression in rice

To investigate if enhanced metabolism through JA-Ile impacts hormone signaling as was reported in Arabidopsis and wild tobacco (Caarls *et al*., 2017, Marquis *et al*., 2022, Smirnova *et al*., 2017, Tang *et al*., 2020), expression of established JA-response markers was quantified in unstimulated 7 d-old rice seedlings of the 8 JAO-deficient lines (Fig. 5). *JAZ9*, *TPS30*, *RBBII-2* and *NOMT* genes were significantly upregulated in most mutant lines. *OsTPS30* is the rice homolog of maize *ZmTPS10* gene involved in the synthesis of the volatile terpene pheromones (E)-β-farnesene and (E)-α-bergamotene (Schnee et al., 2006). *RBBI-2* is JA-regulated proteinase inhibitor with reported antifungal activity (Qu et al., 2003). NOMT is the methyl-transferase catalyzing the terminal biosynthetic step leading to the flavonoid phytoalexin sakuranetin, that exhibits antifungal and anti-herbivore properties (Liu et al., 2023). Of note, their expression was not always reflecting the number of mutated *JAO* genes and the change in hormone levels. Maximal transcript levels were achieved generally in triple mutants while *jao1.2* and *jao1.2.3.4* lines had an intermediate gain in expression. We conclude from this behavior that genetic removal of JA hydroxylation triggers increased defense signaling but that interaction of *jao* mutations results in complex outputs on target gene expression.

**Figure 5.**
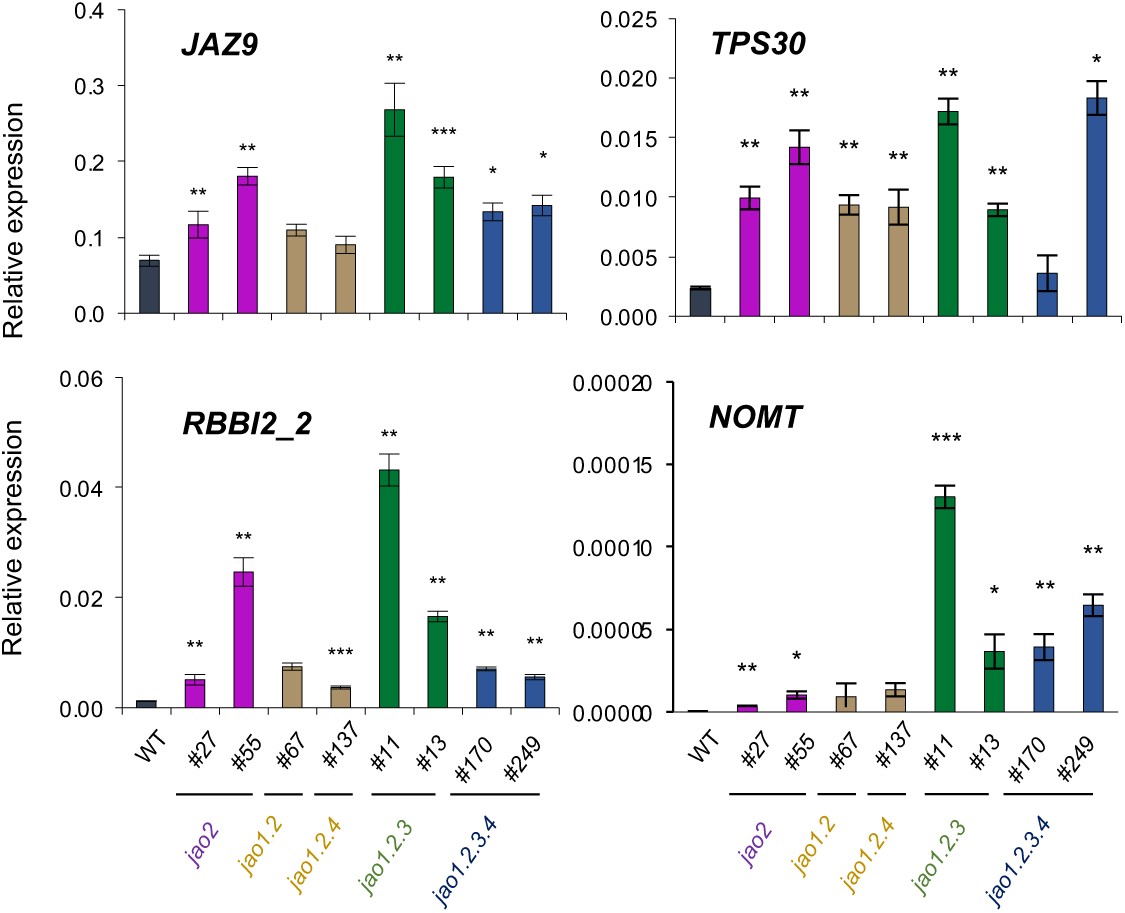
Impact of *jao* mutations on transcript levels of JA-responsive genes in leaves comparatively to WT. Same material as in Figure 3 was used for RNA extraction and RT-qPCR analysis. Expression was normalized with signal from *UBQ5* and *UBQ10* housekeeping genes. Histograms represent means of 4 biological replicates with SEM. Asterisks represent statistical significance for each mutant line compared with WT (t-test, \**P*<0.05; \*\**P*<0.01; \*\*\**P*<0.001).

### Rice *jao* mutations gradually impact leaf metabolic profiles

To get a broader picture of the changes in *jao* mutants, we established a global metabolic profile of young leaves of eight JAO-impaired plant genotypes. Overall distribution of signals in a principal component analysis (PCA) revealed that mutants separate into distinct groups with single *jao2* lines being closest to WT (Fig. 6A). Double *jao1.2* and triple *1.2.4* mutants on the one hand and triple *jao1.2.3* and quadruple *jao1.2.3.4* mutants on the other hand formed additional groups that diverged further from WT. This global assessment indicates an incremental impact of additive *jao* mutations on the medium-polar metabolite content of rice leaves. It also suggests tha*t OsJAO4* impairment does not trigger an additional shift when combined to other mutations. We detected 1849 metabolic features in the sample set (Supplementary Table S1) and their relative abundance was submitted to pairwise comparisons between WT and each individual mutant line as shown in Suppplementary Table S2. From this latter analysis, the number of differential features (down or up) in each mutant are listed in Fig. 6B. It appears that in all lines, the number of down-regulated compounds exceeds the number of up-regulated ones and that the total number or differentials increases with the number of *jao* genes mutated up to triple mutants. When the features were organized in a hierarchical heatmap, contrasted patterns were revealed among genetic groups (Fig. 6C). For example, a set of features (cluster A) was present in WT extracts and their abundance gradually decreased from single *jao2* lines to higher order mutants. Conversely, cluster B groups features of low relative abundance in WT but whose occurrence steadily increased in single to multiple mutants. These findings indicate that *jao* mutations gradually reconfigure sectors of the metabolome in rice leaves with either enhancing or repressing effects on specific sets of compounds. In an attempt to identify some of the JAO-modulated compounds, we interrogated multiple metabolomic databases and established a list of putatively annotated targets (Fig. 6C right panel). This list spanned compounds from several biochemical pathways including indole derivatives, phenylpropanoids, phenolamides or primary metabolites. Of note, some compounds increasing in *jao* mutants were previously reported to display defensive properties. Among overaccumulated compounds are sinapic acid and sinapoylputrescine, belonging to a wound-induced phenolamide class previously described in rice (Tanabe et al., 2016); palmitoylputrescine, displaying antibiotic properties (Brady and Clardy, 2004) ; benzoic acid and hydroxylated derivatives that are fungistatic and antibacterial (Nehela et al., 2021; Zhang et al., 2022); diacetyl-3,6-diferuloylsucrose, known as smilaside M is a potent fungitoxic compound (Zhou et al., 2019). Finally, 5-hydroxyindole-3-acetic acid is a known catabolite of serotonin in animals (Corcuff et al., 2017) and genetic serotonin suppression in rice triggers resistance to destructive insect pests (Lu et al., 2018).

**Figure 6.**
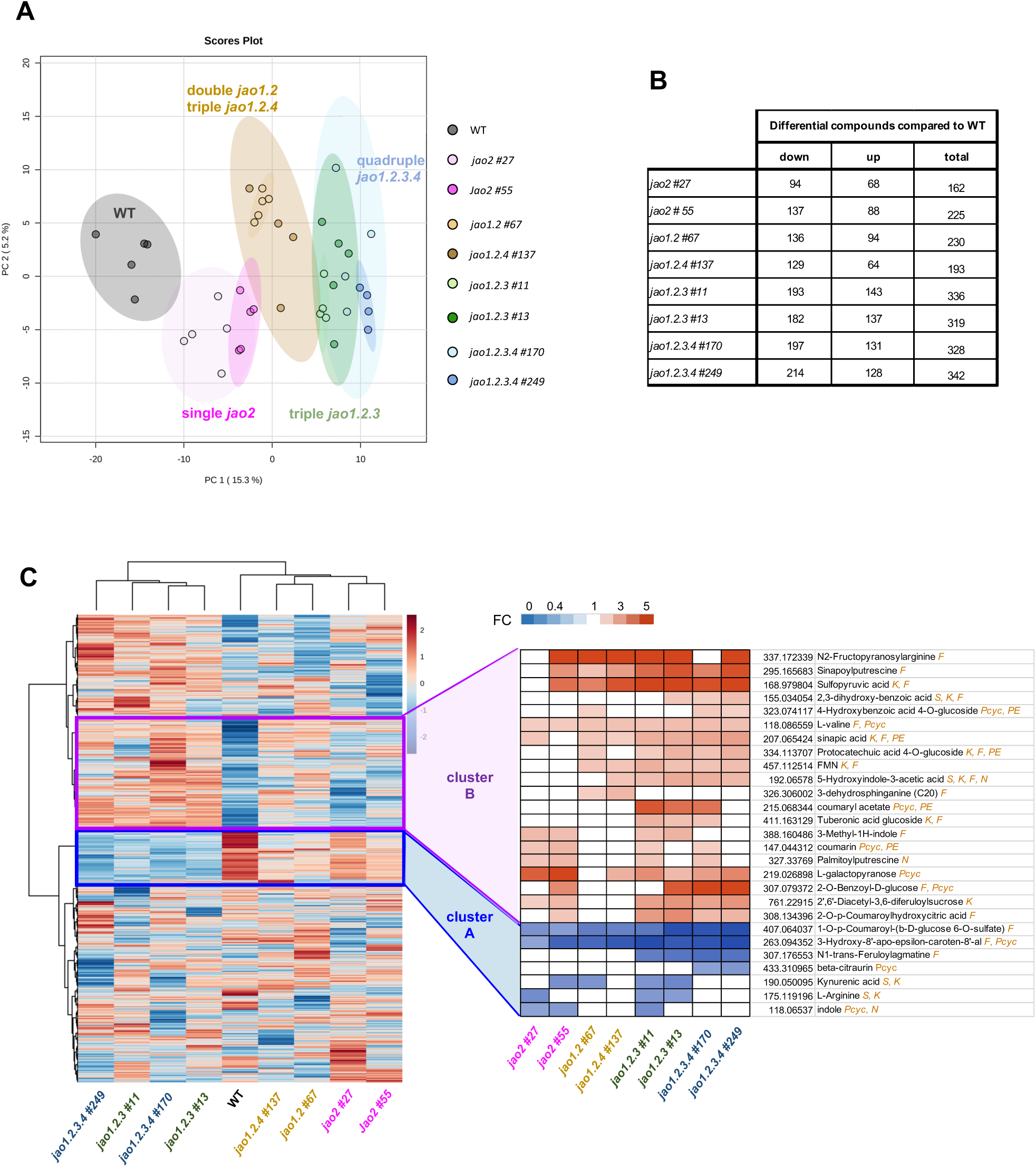
Comparative analysis of non-targeted metabolic profile in WT and 8 mutant lines (from single *jao2* to quadruple *jao1.2.3.4*) with impaired expression of *OsJAO* genes. Leaf material (same as in Figure 3) was extracted and methanol-soluble metabolic content was analyzed by high-resolution LC-MS/MS. (a) Principal Component Analysis (PCA) of the distribution of the metabolomes of 9 genotypes using five biological replicates per genotype. All genotypes had 5 biological replicates except *jao1.2.3 #11*, *jao1.2.4 #137*, *jao1.2.3.4 #170* and *jao1.2.3.4 #249* which had 4 biological replicates. (b) Number of differential compounds between WT and each indicated mutant genotype among 1849 total features detected (Supplementary Table S1; Supplementary Table S2). Statistical analysis was conducted with Metaboanalyst (https://www.metaboanalyst.ca) tool (−1<log_2_FC>1; p-value<0.1). (c) Left panel: hierarchical heatmap established with Metaboanalyst 5.0 representing the comparative evolution in abundance of metabolic features in 8 JAO-impaired mutant genotypes relative to WT. Right panel: close-up of annotated features ranked by global quantitative evolution relative to WT. Values on right represent measured m/z values. Databases producing annotations: F: FoodDB; K: KNApSAcK; N: NPA; PC: PlantCyc; PE: PhenolExplorer; S: Multiple spectral libraries, see Methods section. S symbol elevates annotation confidence to level 2a as defined by Schymanski et al (2014). FC: Fold Change.

### Rice *jao* mutations confer increased basal immunity to *M. oryzae*

To investigate whether the loss of *OsJAO* genes influences immunity to a pathogen, we analyzed the interaction of *jao-2* and *jao1.2.3.4* mutant plants with the fungal rice pathogen *Magnaporthe oryzae*. We performed standard whole plant spray inoculation assays (Berruyer et al., 2003) with the *M. oryzae* isolate Guy11 (*Mo* Guy11). As previously reported, *Mo* Guy11 caused typical blast disease lesions on the leaves of Kitaake WT plants (Fig. 7A, Supplementary Fig. S9A). These lesions were characterized by a dark, brown margin, which is a hallmark of basal immunity, and a grey center, where the fungus sporulates under favorable conditions. In addition to these susceptible lesions, there were also numerous smaller, dark brown lesions, lacking the grey center and that are typical for partial resistance. On the very susceptible variety Maratelli that we used as a positive control, only susceptible type disease lesions were observed (Supplementary Fig. S9A). In the *jao1.2.3.4 #170* mutant, the number of lesions and the leaf area covered by lesions were significantly and strongly reduced as compared to WT plants (Fig. 7A). In the *jao2 #27* mutant, both parameters were also reduced, but to a lesser extent and the differences were not significant. However, the size of the individual lesions was not altered in both mutants as compared to the WT (Supplementary Fig. S9B left panel). These results suggest that the loss of the *OsJAO* genes leads to increased basal immunity in rice. A reduced lesion number without an alteration of the lesion size suggests that *OsJAO* impairment affected primarily the initial invasion of rice epidermal cells by the blast fungus but not the subsequent development of the fungus inside the leaves.

**Figure 7.**
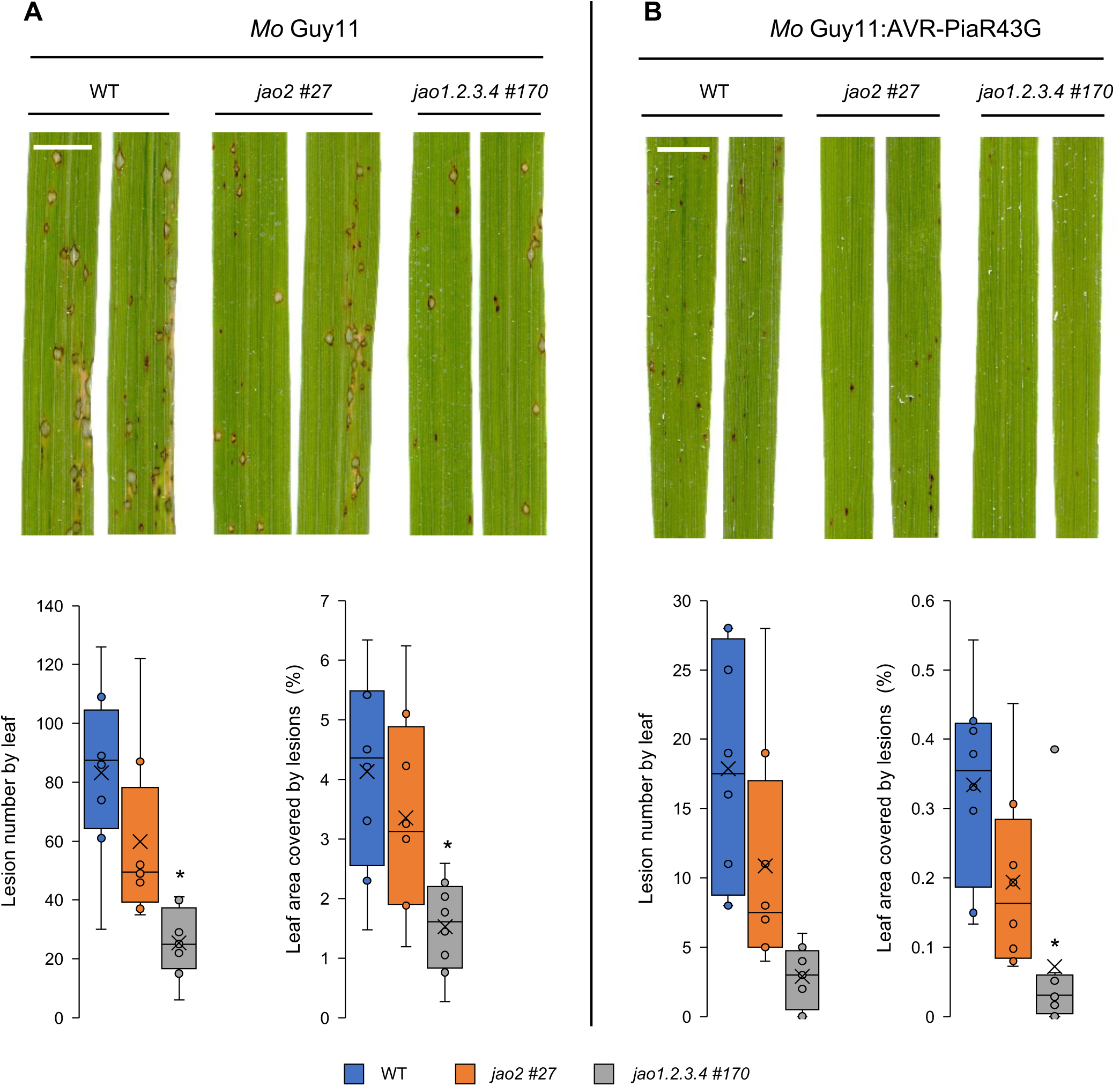
Loss of *OsJAO* genes results in increased basal resistance to *M. oryzae*. Plants of the Kitaake variety (WT and of *jao-2 #27* and *jao1.2.3.4 #170* mutants) were spray-inoculated with spores of the *M. oryzae* strain (A) Guy11 carrying an empty vector (*Mo* Guy11) or (B) a Guy11 strain possessing AVR-PiaR43G (*Mo* Guy11:AVR-PiaR43G). Images were taken by scanning of leaves 7 days after inoculation. Pictures show representative symptoms (upper panel). Scalebar: 1 cm. Plots in lower panels show numbers of lesions per leaf (n=8) and leaf area covered by lesions as determined by image analysis. The boxes represent the second quartile, median, and third quartile. Individual data points are presented as circles and mean values as crosses. The mean values of the *jao1.2.3.4 #137* mutant were according to a t-test significantly different from those of the Kitaake WT with p<0,005. They are therefore marked with *. The mean values of the *jao2 #27* mutant were not different. Equivalent results were obtained in two other independent inoculation experiments.

To investigate whether the loss of the *OsJAO* genes influences specific, NLR-mediated resistance, we inoculated the *jao* mutants with an *Mo* Guy11 strain expressing the effector AVR-Pia (Guy11:AVR-Pia). AVR-Pia is recognized in Kitaake by the CNL sensor/helper pair Resistance Gene Analog 4 and 5 (RGA4/RGA5) triggering hypersensitive response (HR) and preventing *Mo* infection (Cesari et al., 2013). The *Mo* Guy11:AVR-Pia strain caused no visible symptoms or very small HR lesions on Kitaake WT and *jao* mutant plants, and no difference in the resistance level was observed between the three rice lines (Supplementary Fig. S9C).

To test whether weaker NLR-mediated immune activation would reveal alterations of immunity in the *jao* mutants, we performed inoculation experiments with an *Mo* Guy11 strain expressing an AVR-Pia variant with attenuated RGA5 binding due to the replacement of an arginine residue at position 43 by a glycine (Ortiz et al., 2017). This *Mo* Guy11:AVR-PiaR43G strain triggered weakened resistance on Kitaake characterized by more and larger HR lesions in comparison to the *Mo* Guy11:AVR-Pia wild-type isolate (Fig. 7B, Supplementary Fig. S9D). These extended brown lesions caused by the *Mo* Guy11:AVR-PiaR43G strain on Kitaake leaves were reminiscent of partial resistance. On *jao1.2.3.4 #170* mutant plants, the *Mo* Guy11:AVR-PiaR43G strain caused less lesions and a smaller total lesion area than on WT Kitaake (Fig. 7B). In addition, the area of individual lesions was significantly reduced (Supplementary Fig. S9B). On leaves of the *jao2 #27* mutant, the number of lesions and the total lesion area per leaf was also reduced, but to a lesser extent than on the quadruple mutant and the differences were again not significant.

We conclude from these experiments that NLR-triggered immunity is not altered in *jao* mutant plants. However, loss of *JAO* gene function resulted in reduced lesion numbers in compatible or partially resistant interactions, without a change in lesion size in the compatible interaction. This finding indicates that the loss of JAO functions limits the frequency of successful fungal penetrations but does not alter the subsequent development of the fungus inside the plant leaf. This phenotype could be a consequence of a constitutive activation of defense responses in *jao* mutant plants.

To link the phenotypes in the interaction with *Mo* Guy11:AVR-PiaR43G with defense activation in the three plant genotypes, we monitored the expression of representative markers of various JA-regulated defense pathways at 2 days post-inoculation (dpi), when the outcome of the interaction is determined. We could not evidence, in any gene examined, a difference in expression amplitude in infected tissue in mutants relative to WT (Supplementary Fig. S9E). In contrast, expression of the repressor *JAZ8*, the indole biosynthetic gene *IGL* (Frey et al., 2000), or the diterpenoid phytoalexin biosynthetic gene *TPS29* (Sun et al., 2022) were higher in uninfected mutant plants. This corroborates that *osjao* mutations enhance defensive capacities of leaves prior to infection.

## Discussion

2-ODDs are involved in virtually every plant hormone metabolic pathway - including abscisic acid, ethylene, gibberellins, salicylic acid, or strigolactones -, either in biosynthetic or catabolic processes. In the JAs pathway, recently-characterized JA oxidases have a peculiar position because they define a metabolic diversion that dampens the activation of JA into JA-Ile. Although several other JA-modifying enzymes are known (Wasternack and Strnad, 2018), *jao* genetic impairment has revealed its unique function in attenuating basal JA-Ile signaling in Arabidopsis leaves (Smirnova et al., 2017). Transient silencing of homologous genes boosted direct and indirect defenses in the wild tobacco *N. attenuata* (Tang et al., 2020). In both species, impaired JAO function was associated with enhanced tolerance to fungal or insect attacks, as well as better survival to severe drought in Arabidopsis (Caarls et al., 2017; Smirnova et al., 2017; Tang et al., 2020; Marquis et al., 2022). Interestingly, these gains in defensive capabilities were observed without major growth penalty except in the Arabidopsis quadruple *jaoQ* mutant. The *jao* pathway therefore bears promising features as to modulate the strength of JA-regulated responses, and its potential conservation and impact needs to be investigated in crop species.

Here we addressed the occurrence, activity and functions of the JAO pathway in rice, a monocot of uppermost agronomic importance. Four predicted homologs of the regulatory AtJAO2 protein were readily identified in rice genome for functional analysis. Their sequences cluster in a monocotyledon clade along with similar proteins from maize, sorghum or barley, OsJAO1, 2, and 3 being closest to characterized dicot JAO proteins. OsJAO4 positions further away in a cereal-specific subclade, at the junction with a wide group of proteins acting in flavonoid metabolism. All OsJAO obtained as recombinant proteins but OsJAO4 readily catalyzed oxidation of JA to 12OH-JA, and these enzymes reverted the high basal defense phenotype when ectopically expressed in the Arabidopsis *jao2-2* mutant. This demonstrates that the biochemical function of JAO2 is conserved in rice OsJAO1/2/3 proteins and that they attenuate JA-regulated defense responses. The differential responsiveness of *OsJAO1*, *2 and 3* expression to MeJA exposure indicates that each isoform is recruited at distinct timescales, and calls for determining their specific or overlapping sites of expression throughout rice development and adaptation. Despite sharing with the other OsJAOs conserved residues critical for JA substrate binding (Zhang et al., 2021), no consistent functional data could be obtained for OsJAO4. We hypothesize that this protein is either inactive under our conditions or performs a different reaction that may be restricted to roots, as no expression could be detected in leaves.

The use of a multiplex Crispr/Cas9 mutagenic construct allowed to readily obtain an original series of modified plant lines, including double, triple and quadruple mutants, providing access to the consequences of incremental suppression of JAO activity. The examination of these plant genotypes has revealed several phenotypes reminiscent of enhanced JA signaling. The primary observed effect of reduced OsJAO activity was the redirected metabolic flux towards JA-Ile and its catabolites. A mild but significant impact was recorded in single *jao2* alleles, that remained stable when *jao1* or *jao4* mutations were added to *jao2*, suggesting a minor contribution of OsJAO1 and OsJAO4 to leaf JA homeostasis. This will need the analysis of *jao1.2* alleles to be confirmed in future studies. In contrast, combined *jao2.3* mutations, either in two triple *jao1.2.3*, or in two quadruple *jao1.2.3.4* lines, resulted in a stronger metabolic shift, demonstrating functional redundancy of OsJAO2 and OsJAO3 in shaping the basal JAs profile in rice leaves under stress-free conditions. This gradual flux redirection effectively upregulates *JAZ* repressor and basal defense gene expression, a hallmark of higher constitutive JA signaling, but their fluctuating expression patterns through the mutant series suggest that factors additional to steady-state hormone levels determine the responsiveness of target genes.

The broader biochemical consequences of redirected JA/JA-Ile catabolism were addressed by non-targeted metabolomic exploration of the same nine genotypes. PC analysis disclosed a remarkable gradual impact of *jao* impairment on global leaf metabolome and additionally detected an intermediate status of *jao1.2* alleles that was not predicted by JAs profiles. This picture highlights the potential of modulating JAO activity to fine-tune quantitatively JA responses in rice. Of note, when differential abundance of annotated compounds was filtered out relative to WT, typical defense metabolites of the genus *Oryzae* did not show up, as was the case for *Brassicaceae-*specific glucosinolate metabolites in the Arabidopsis *jao2* line (Marquis et al., 2022). Instead, elevated contents in phenylpropanoid, phenolamide, indole, or hexose sugar derivatives were recorded in rice, of which several were reported as JA-regulated or bearing antimicrobial properties. This illustrates that the JAO pathway impacts branches in both primary and specialized metabolism.

Consistent with the pre-existing accumulation of JA-regulated defense proteins and metabolites, *jao2* and particularly *jao1.2.3.4* mutants were more resistant to both virulent and avirulent strains of *M. oryzae*. The important role of JA in the basal resistance of rice to *M. oryzae* is well documented (Nguyen et al., 2019), and it has been shown that temperature-dependent variations in JA signaling strongly influence the basal immunity of rice to the blast fungus (Qiu et al., 2022). Consistent with this, the blast fungus deploys virulence functions that interfere with JA signaling in the host plant (Patkar et al., 2015). In our study, we show that constitutively increased levels of JA-Ile due to impaired catabolism of the hormone precursor provides strong protection of rice against *M. oryzae*. However, the induction of immunity responses upon detection of the fungus and limiting its *in planta* development seem not or only marginally modified in the *jao* mutant plants. This is consistent with a role of *OsJAO* genes in modulating JA levels and signaling mostly in unstressed plants, with possible consequences on the plant holobiont (Carvalhais et al., 2017). Further studies will be needed to examine the interaction of JAOs with other hormonal pathways (Lahari et al., 2024) or their impact on tolerance to various biotic stresses in rice.

Examination of growth and fitness parameters in seedlings of JAO-impaired lines highlighted interesting features and provides a readout of the importance of intricate JA metabolism on growth/defense tradeoff. While the development of the first leaves was particularly affected by total JAO deficiency (*jao1.2.3.4*), negative effects on other growth parameters was milder and comparable in the different mutant lines. Interestingly, the vegetative growth penalty was attenuated at later developmental stages, indicating that JAO activity is required mostly at early seedling development to optimize growth in rice. In addition, the reduced seed mass of *jao* mutants points to the need to fine-control the extent of JA signaling to execute proper reproductive programs. It remains to be determined if this outcome results from a reduced supply of photosynthates to the panicle, or if excessive JA signaling impairs flower-specific processes.

In conclusion, we demonstrated that the JAO pathway is biochemically conserved between monocot and dicot species, and most importantly is also endowed with a regulatory function in rice, at the nexus of growth/defense balance. By diverting metabolic flux upstream of active JA-Ile formation, JAO activity optimizes rice juvenile growth, maximizes seed yield and attenuates the expression of baseline defense programs. While significant redundancy was evidenced in leaves, it is possible that individual OsJAO isoforms exert more specific repression in discrete organs/tissues or under particular situations. Their identification could provide tools to tailor adequate levels of JA signaling to adapt growth/defense balance to meet specific agronomical traits. Similarly to oxidative JA-Ile catabolism whose natural variation has contributed to rice adaptation to a temperate climate (Mao et al., 2019), varying levels of JAO activity may be a determinant of JA signaling activity within the wide rice genetic diversity.

## Material and methods

### Rice cultivation

*Oryza sativa* L. ssp. japonica cv. Kitaake was used throughout the study. Seeds were dehusked before being sterilized as described in Ndecky et al (2023). Seeds were then sown in Magenta boxes containing 0.4 % phytoagar (358 mg l^-1^ basal Murashige and Skoog (MS) solution (Duchefa Biochemie) buffered to pH 5.8 with MES. After 7 days *in vitro* culture in a growth cabinet (Percival Select 41L, Perry, Iowa, USA) under a 12h/12h photoperiod with 125 µmol light at 28°C, the seedlings were transferred to soil and incubated under the same conditions. The growth-related phenotyping was performed after 3 weeks growth on soil in a greenhouse under a 16h-day/8h-night photoperiod, at 28°C/24°C with a 75% relative humidity.

### Magnaporthe oryzae (Mo) infection assays

For inoculation experiments, rice plants were grown in soil in a growth chamber at 28°C, 16 h light and 65% relative humidity. *Magnaporthe oryzae* (Mo) strains Guy11, Guy11 transformed with AVR-Pia or AVR-Pia_R43G (Ortiz et al., 2017) were grown as described (Faivre-Rampant et al., 2008). *Mo* conidial spores were collected frorm eight to ten old cultures in water with 0.5 % gelatine, Suspensions of conidia were adjusted to a concentration of 5 × 10^4^ spores ml^−1^ and sprayed onto 3-week-old rice plants (Berruyer et al., 2003). Inoculated plants were transferred for 16 h in a dark growth chamber regulated at 26°C and 100% humidity and then placed in normal growth conditions. Symptoms were recorded seven days after inoculation by imaging leaves and measuring lesions and corresponding leaf areas using the in-house developed program ‘Leaf Tool’ (Cesari *et al*., 2022).

### Arabidopsis cultivation

All *Arabidopsis thaliana* genotypes used in this study were in a Col-0 ecotype. *atjao2-2* (GK_870C04) T-DNA line was obtained from the Nottingham Arabidopsis Stock Center (NASC, https://arabidopsis.info). Individual ORFs of *OsJAOs* genes were inserted into the pEAQΔP19 plasmid. Recombinant plasmids were mobilized in *Agrobacterium tumefaciens* GV3101 strain before transforming the *jao2-2* line with the floral dip method (Clough and Bent, 1998). Primary transformants (T1) were selected on kanamycin-containing MS plates. Kanamycin-resistant seedlings were transferred to soil under a 16h/8h day/night photoperiod (21°C/16°C) in a growth chamber. After two weeks cultivation, unstimulated leaves were collected from 15-20 individual T1 transformants for RNA extraction and defense complementation assay by RT-qPCR. Two independent single copy, complementing lines were conducted to homozygotes in T3 generation.

### Assembly of plasmid constructs

#### Plasmids for JAOs heterologous expression in Arabidopsis

The gene constructs used to overexpress rice OsJAO proteins in the *atjao2-2* Arabidopsis mutant were assembled with the Golden Gate (GG) method into the plasmid pEAQΔP19-GG, a derivative of the pEAQ-HT plasmid lacking the P19 gene (Incarbone *et al*., 2021). The T-DNA of the pEAQΔP19-GG plasmid has two *Sap*I restriction sites located downstream of a 35S promoter (p35S). The coding sequence (ORF) of each individual OsJAO (wild type or mutated) ORF was inserted downstream to the Green Fluorescent Protein (eGFP) ORF. To this end, each *OsJAO* ORF was amplified by PCR from cDNAs produced by retrotranscription of rice plant mRNAs. Forward primer covered about 20 nucleotides upstream of the initiator codon and the reverse primer covered about 20 nucleotides (nt) 3’ of the stop codon. The *Sap*I restriction sites were grafted to the 5’ and 3’ ends of these amplicons through a second PCR. Each OsJAO module was assembled with the eGFP module in the pEAQΔP19-GG plasmid using Golden Gate while ensuring the assembly maintains OsJAO sequences always downstream of eGFP sequence and both OsJAO and eGFP encoded in a unique heterologous CDS. The resulting plasmids were transformed into *E. coli* Top10 strain, selected then validated by Sanger sequencing.

### Plasmids for bacterial expression

The gene constructs used to express the recombinant proteins 6xHis-MBP-OsJAO4 were assembled by GG into the pETGG plasmid, a Golden Gate derivative from pET22b(+) plasmid. with two *SapI* restriction sites, downstream of the lac operator.

The construct used to express the recombinant 6xHis-MBP-OsJAO2 protein was assembled by Gateway (GW) cloning into the pHMGWA plasmid (Busso et al., 2005) using pDONRzeo as an intermediate. The plasmid constructs used to express the recombinant 6xHis-OsJAO1 and 6xHis-OsJAO3 protein were also assembled into the pHGWA (Busso et al., 2005) plasmid by GW method.

### Plasmids for rice mutagenesis

#### General strategy

Rice mutagenesis was performed using CRISPR-Cas9 strategy thanks to two simplex constructs pUbi-Cas9:sgRNAs-OsJAO1 and pUbi-Cas9:sgRNAs-OsJAO2 used to generate *jao1* and *jao2* single mutants respectively. These two plasmids encode mainly the Cas9 enzyme and two sgRNAs specific to the targeted gene. A third plasmid pUbi-Cas9:sgRNAs-Multiplex whose T-DNA encodes one sgRNA for each of the 4 *JAO* genes was used to produce different variants of multiple mutants. All three recombined plasmids were assembled using the pUbi-Cas9 and pENTR4:gRNA4 vectors based on the methods reported by Zhou et al. (2014) and Xie et al (2015).

All the primers used in the study are listed in Supplementary Table S3. The CRISPR RNAs (crRNA) crRNA8-OsJAO1 and crRNA9-OsJAO1 were used to target the *OsJAO1* gene. crRNA9-OsJAO1 was first assembled by hybridization of the 24-nt long primers crRNA9-OsJAO1_fw and crRNA9-OsJAO1_rv that are complementary on the 20 nt of their 3’ end that correspond to the sequence of crRNA9-JAO1. The 4 nt in the 5’ of these primers are complementary to the sticky ends generated by the cleavage of pENTR4:gRNA4 by the *Bsa*I enzyme. By insertion between the *Bsa*I sites, crRNA9-JAO1 was placed under the control of the U6p2 promoter and forms, together with the trans-activating crRNA (tracrRNA) downstream of the promoter, sgRNA9-OsJAO1. The pENTR4:sgRNA9-OsJAO1 plasmid formed was subsequently used to insert crRNA8-OsJAO1 between the *Btg*ZI sites. crRNA8-OsJAO1 was assembled in the same way as crRNA9, by hybridization of the complementary primers crRNA8-OsJAO1_fw and crRNA8-OsJAO1_rv. The 4 nucleotides in the 5’ of the latter two primers are complementary to the sticky ends generated by the cleavage of pENTR4:sgRNA9-OsJAO1 by the *Btg*ZI enzyme. Thanks to its insertion between the *Btg*ZI sites, crRNA8-OsJOA1 was placed under the control of the U6p1 promoter and forms, together with the tracrRNA downstream of the promoter, sgRNA8. The plasmid pENTR4:sgRNA9-OsJAO1 thus became pENTR4:sgRNAs-OsJAO1. For yet unknown reasons, recombining pENTR4:sgRNAs-OsJAO1 with pUbi-Cas9 by LR recombination was impossible. As an alternative, we PCR-amplified the assembled construct of the pENTR4:sgRNAs-OsJAO1 plasmid and transferred it into the pENTR1A entry plasmid using BP recombination. LR recombination of the neoformed pENTR1A:sgRNAs-OsJAO1 with the pUbi Cas9 plasmid generated the vector pUbi-Cas9:sgRNAs-OsJAO1.

crRNA1-OsJAO2 and crRNA4-OsJAO2 were used to target the *OsJAO2* gene. We used the same method as that used to assemble the pUbi-Cas9:sgRNAsOsJAO1 plasmid. In this case, crRNA1-OsJAO2 was placed under the control of the U6p1 promoter and crRNA4-OsJAO2 under the control of the U6p2 promoter.

For the multiplex construct, a single crRNA was used to target each of the 4 *OsJAOs* genes. The *OsJAO1* gene was targeted by crRNA9-OsJAO1, the *OsJAO2* gene by crRNA4-OsJAO2, the *OsJAO3* and *OsJAO4* genes were targeted by crRNA4-OsJAO3 and crRNA1-OsJAO4, respectively (Supplementary Table S3). To facilitate the assembly of the sgRNAs, we reused the pENTR4:sgRNA9-OsJAO1 plasmid in which sgRNA9-OsJAO1 is already assembled in front of the U6p2 promoter. The sgRNAs of the *OsJAO2*, *OsJAO3*, and *OsJAO4* genes were assembled in this plasmid in front of the U6p1 promoter as a polycistronic sequence according to a multiplexing method reported by Xie et al. (2015). Because the assembled plasmid pENTR4:sgRNAs-Multiplex was unable to recombine with pUbi-Cas9 through an LR reaction, the generated sgRNAs construct was PCR-amplified was transferred into the pENTR1a entry plasmid. LR recombination of the neoformed pENTR1a-sgRNAs-Multiplex with pUbi-Cas9 produced the pUbi-Cas9-sgRNAs-Multiplex plasmid used for the production of multiple mutants.

### Plant transformation

#### Production of Arabidopsis transgenic lines

Recombinant plasmids were mobilized into *Agrobacterium tumefaciens* GV3101 strain before transforming the *jao2-2* line with the floral dip method (Clough and Bent, 1998). Primary transformants (T1) were selected on kanamycin-containing MS plates. Kanamycin-resistant seedlings were transferred to soil under a 16h/8h photoperiod (21°C/16°C) in a growth chamber. After two weeks cultivation, unstimulated leaves were collected from 15-20 individual T1 transformants for RNA extraction and defense complementation assay by RT-qPCR as described in Smirnova *et al*. (2017). Two independent single-copy, complementing lines were conducted to homozygotes in T3 generation for each *OsJAO* gene.

#### Production of rice knock-out mutants

Genetic transformation of rice (*O. sativa*, cv. kitaake) was performed on embryogenic calli based on the protocol described by Hiei and Komari (2008). Briefly, *A. tumefaciens* bacteria (strain EHA105) transformed with the plasmid constructs pUbi-Cas9:sgRNAs-OsJAO1 or pUbi-Cas9:sgRNAs-Multiplex were co-cultured with rice calli to induce T-DNA transfer into the genome of callus cells. The newly transformed callus cells were then selected based on their resistance to hygromycin conferred by the *HPTII* gene present on the T-DNA and regenerated into seedlings to constitute the T0 generation of transgenic plants. Genomic DNA was isolated from leaves of T0 plants to confirm the presence of a T-DNA by PCR. Mutations in target genes were then searched for using two analytical methods. First, the plants of interest were subjected to High Resolution Melting (HRM) analysis on a LightCycler 480 II instrument (Roche Applied Science, Penzberg, Germany) to identify individuals with mutations in the target genes. The analysis was performed in a 10 µL reaction with Precision Melt Supermix kit (Bio-Rad) according to manufacturer’s instructions. When positive, the nature of the induced mutations was revealed by Sanger sequencing. When the mutation was heterozygous, an analysis of the chromatograms generated by sequencing with the ICE-Synthego tool (Conant *et al*., 2022) was necessary to distinguish the different alleles present at the locus concerned. Next, in the T1 and T2 progeny of these plant lines, plants that were devoid of T-DNA (Cas9-free) and knock-out (ko) for the targeted *OsJAOs* genes were further identified by PCR.

### Expression and Purification of recombinant proteins

*Escherichia coli* bacteria of the Rosetta 2 pLyS strain transformed with pHGWA:OsJAO1, pHMGWA:OsJAO2, pHGWA:OsJAO3 and pETGG:6xHis-HMP-OsJAO4 were first cultivated at 37°C and 250 rpm in LB medium to reach the optical density (A_600_) of 0.45 and 0.5. Expression of the recombinant proteins was induced by adding IPTG 0.5 mM and the cultures further incubated at 20° C for 4 h. Cells were then collected by centrifugation (20 min, 3000 g, 4° C) and the bacterial pellets frozen at −80°C until processing. Recombinant protein expression was analyzed in total bacteria by western blot using a monoclonal mouse anti-6x-his antibody (Covalab, Bron, France) and goat anti-mouse secondary antibody coupled to horseradish peroxidase (Invitrogen). Signal was recorded using luminescence on a Fusion Fx instrument (Vilber Lourmat, Marne-la-Vallée, France). For JAO activity assays to be performed on purified proteins, the thawed bacterial pellets were resuspended in lysis buffer (300 mM NaCl, 20 mM imidazole in 50 mM Tris-HCl pH 7.5) at 20 A_600_ units mL^-1^. Bacteria were lysed by sonication (VibraCell™ 75115, Bioblock Scientific) and cell debris removed by centrifugation (15 min, 17000 g, 4°C). The clarified lysate was filtered at 0.22 µm and the proteins of interest purified by affinity chromatography by affinity chromatography on an Äkta pure Fast Protein Liquid Chromatography system equipped with a Histrap FF crude 1ml IMAC (immobilized metal affinity chromatography) column (Cytiva). Equilibration and binding and washes were done with 50 mM Tris pH 8, 300 mM NaCl, 5% glycerol, 25 mM imidazole and elution with 50 mM Tris pH 8, 300 mM NaCl, 5% glycerol and 500 mM imidazole. Eluted fractions were analysed by Coomassie Blue-stained polyacylamide gel or by Western blot. The protein concentration of eluates was estimated by quantification with the Bradford reagent using a bovine serum albumin (BSA) calibration range. For the enzymatic tests performed directly with the bacterial lysates, the collected cell pellets were resuspended in buffer devoid of imidazole (300 mM NaCl, 5 mM dithiothreitol (DTT) in 50 mM TrisHCl pH 7.5), and cells lyzed by sonication (VibraCell™ 75115, Bioblock Scientific). Cell debris was then removed by centrifugation (15 min, 17000 g, 4° C) and clarified lysates used for enzymatic assay.

### *In vitro* JA oxidation assay

#### Assays with purified enzymes

Each assay was performed with 10 µg purified OsJAO protein in 200 µL reactions consisting of 100 µM JA substrate, 50 mM Tris-HCl pH 7.5, 5 mM DTT, 13.3 mM 2-oxoglutarate, 13.3 mM ascorbate, 0.67 mM FeSO_4_ and 200 µg BSA. The reactions were incubated at 30° C for 1 h before being stopped with 20% (v/v) HCl 1 M. The reaction products were subsequently extracted with one volume of ethyl acetate. The organic upper phase (200 µL) was transferred in a new tube and dried under nitrogen flow before redissolving in 150 µL methanol. The extracts were analyzed by ultra-high-performance liquid chromatography coupled to tandem mass spectrometry (UPLC-MS/MS) for the detection of JA and its oxidized form OH-JA produced by JAOs as described in Smirnova *et al*. (2017).

#### Assays with clarified bacterial lysates

For OsJAO1 that did not bind properly to affinity resin, assays were conducted with 150 µL bacterial lysate in 300 µL reactions including 30 µM JA substrate, 50 mM Tris HCl pH 7.5, 5 mM DTT, 50 mM 2-OG, 50 mM ascorbate, 0.67 mM FeSO_4_ and 15 µg BSA. The mixtures were incubated at 30°C for 1 h and the reaction products extracted and analyzed following the same procedure used after the enzymatic assays with the purified proteins.

### Extraction of metabolites

Between 35 and 40 mg of frozen fine powder of the leaf plant material were extracted with 80% methanol containing 0.5% acetic acid and 1 µM 9,10-dihydro-JA-Ile, and 100 nM Prostaglandin A1 (Cayman Chemicals, Ann Arbor, Michigan, USA). Material was extracted in 2 ml screw-capped microtubes in the presence of 12 μL mg^-1^ extraction solution by homogenizing with an orbital grinder (Precellys Evolution, Bertin Technologies, France) in the presence of glass beads (2 cycles of 30 s at 6500 rpm separated by 30 s pause). After 1 h of incubation at 4° C on a rotating wheel, the cell debris were sedimented by two successive centrifugations (10 min, 11 000 g, 4° C) and 120 µL of the supernatant were transferred to LC vials containing a glass insert (N9, Machery-Nagel, Düren, Germany).

#### Jasmonate profiling

Jasmonate profiling was performed by LC-MS/MS as described in Marquis et al. (2022) on extracts described above.

#### Metabolome analysis

Untargeted metabolite analysis was performed on same extracts than used for JA profiling on an UltiMate 3000 UHPLC system (Thermo Fischer Scientifc, Illkirch, France) coupled to the ImpactII high resolution Quadrupole Time-of-Flight (QTOF) spectrometer (Bruker, Wissembourg, France). Chromatographic separation was performed on an Acquity UPLC ® HSS T3 column (2.1 × 100 mm, 1.8 µm, Waters) coupled to an Acquity UPLC HSS T3 pre-column (2.1 × 5 mm, 1.8 µm, Waters). The raw data extracted from these analyses were processed with MetaboScape 4.0 software (Bruker): the molecular characteristics were considered and grouped into “buckets” containing one or more adducts and isotopes of the detected ions with their retention time and MS/MS information when available. The parameters used for the definition of the buckets are: a minimum intensity threshold of 5 000, a minimum peak length of 3 spectra, a signal-to-noise ratio (S/N) of 3 and a correlation coefficient threshold of 0.8. The ion [M+H]^+^ was allowed as primary ion, the ions [M+Na]^+^, [M+NH4]^+^ and [M+K]^+^ were allowed as possible seed ions. Biological replicates of the same genotype were grouped together and only the buckets found in 80% of the samples in a group were extracted from the raw data. The resulting list was annotated using SmartFormula to generate a raw formula based on the exact mass of the primary ions and the isotopic pattern. The maximum allowable variation in mass (Δm/z) was set at 3 ppm, and the maximum mSigma value (assessing isotopic model conformity) was set at 30. To name the resulting formulae, analyte lists were derived from FooDB (http://foodb.ca), KNApSacK (http://www.knapsackfamily.com/KNApSAcK/), PlantCyc (https://plantcyc.org/), PhenolExplorer (http://phenol-explorer.eu/), NPA and spectral libraries (Bruker MetaboBASE, Mass Bank, LipidBlast, MSDIAL LipidsDB). The parameters used for the annotation with the analyte lists are the same as for the SmartFormula annotation. The resulting annotations are at the level 3 (analyte lists) and level 2 (spectral libraries) of the Schymanski classification (Schymanski et al., 2014). Statistical analysis was performed with MetaboAnalyst 5.0 (https://www.metabonalyst.ca) with 5 samples per group using peak areas as the reference unit. Data were normalized by sum before log transformation and Pareto scaling. Compounds were considered statistically differential between two groups using thresholds of p-value ≤ 0.1 and fold change ≥ 2 or ≤ −2.

### RNA extraction and gene expression profiling

Leaf samples were ground using the glass-bead homogenizer before RNA isolation using TRizol Reagent according to manufacturer instructions. For RT-qPCR analysis, cDNA was synthesized with Superscript IV Reverse Transcriptase (Thermofisher) using 2 µg of RNA. qPCR was performed using 20 ng of cDNA on a LightCycler 480 II instrument (Roche Applied Science, Penzberg, Germany) as described in Berr *et al*. (2010). The expression levels of the different rice targets genes were normalized against the expression level of the reference genes *UBQ5* (Os01g0328400) and *UBQ10* (Os02g0161900) and the expression level of the Arabidopsis target gene *AtPDF1.2* was normalized against the expression level of the reference genes *AtEXP* (At4g26410) and *AtTIP41* (At4g34270). The sequences of all primers used are listed in Supplemental Table S3.

### Genomic DNA extraction

A small piece of frozen leaf from each rice plant was placed in wells of 96-sample plates and dry-ground in the TissueLyser II (Qiagen) using steel balls. Samples were then ground a second time in the presence of 500 µL Edwards buffer (250 mM Tris HCl pH 7.4, 250 mM NaCl, 25 mM EDTA, 0.5 % SDS), the plate centrifugated (maximum speed, 10 min) to sediment the cell debris. One hundred µL of the supernatant from these cell extracts was recovered and mixed with 75 µL of isopropanol to precipitate DNA. After centrifugation (maximum speed, 15 min), the DNA pellet was washed once with 70% ethanol before being resuspended in 50 µL of distilled water.

### Protein sequence analysis

Protein sequences used in the phylogenetic classification of rice JAOs and their sequence study were extracted were retrieved from the comparative genomics portal Phytozome (https://phytozome next.jgi.doe.gov/). The MEGA-X software (Kumar et al., 2018) was then used to mine protein sequences. The multiple sequence alignment was performed with the Muscle method and the phylogenetic tree drawn with the “Maximum likelihood” method.

## Acknowledgments and Funding

SN was supported by an international doctoral fellowship from the Investissements d’Avenir (IdEx) program from Université de Strasbourg and CNRS. The study benefited from mobility grants from CampusFrance, and a grant from the European Campus (Eucor) program. We thank J. Zumsteg for technical assistance in TQ-LC-MS, T.H. Nguyen for help in rice plant transformation and D. Pflieger for help in data processing.

## Author contributions

TH, SN, TK, AC and MR collectively designed the experiments. SN, TH, LV, NB, CV, VC, IM, DBV performed the experiments. TH, SN, TK, AC and MR analyzed data. TH wrote the manuscript with input of SN, TK, AC and MR.

## Conflict of interest

The authors declare having no conflict of interest.

## Legends to Figures

**Supplemental Figure S1:**
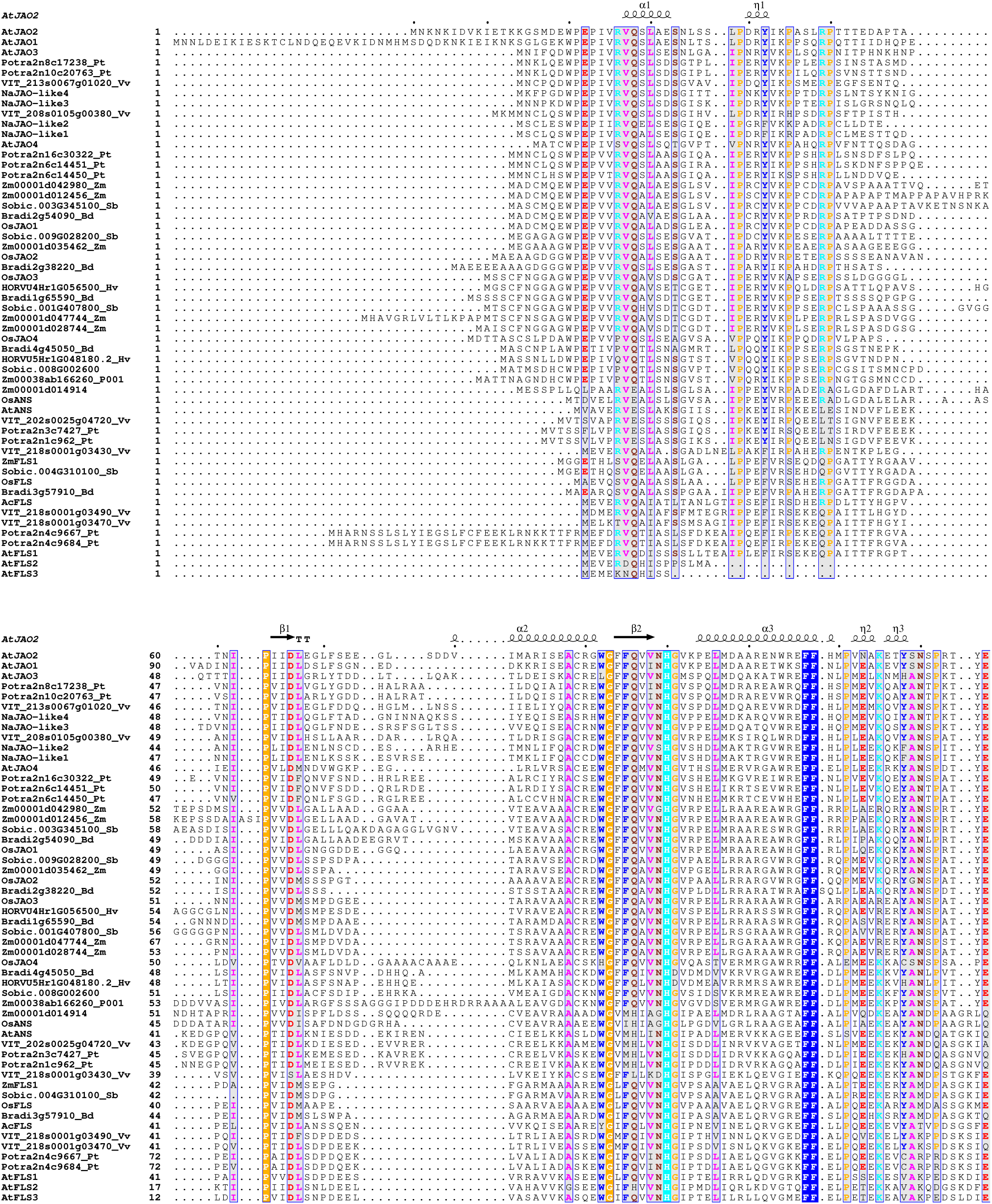
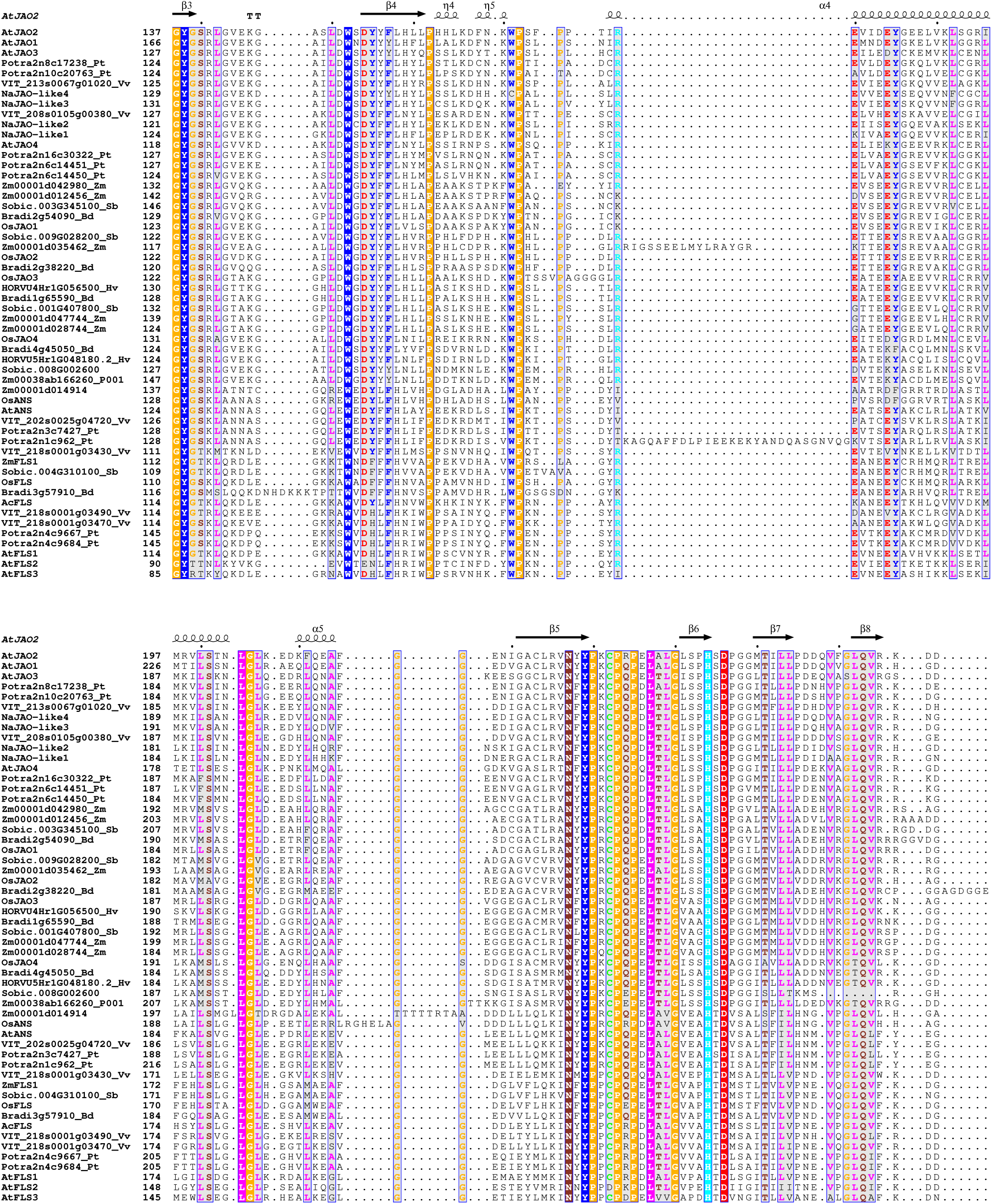
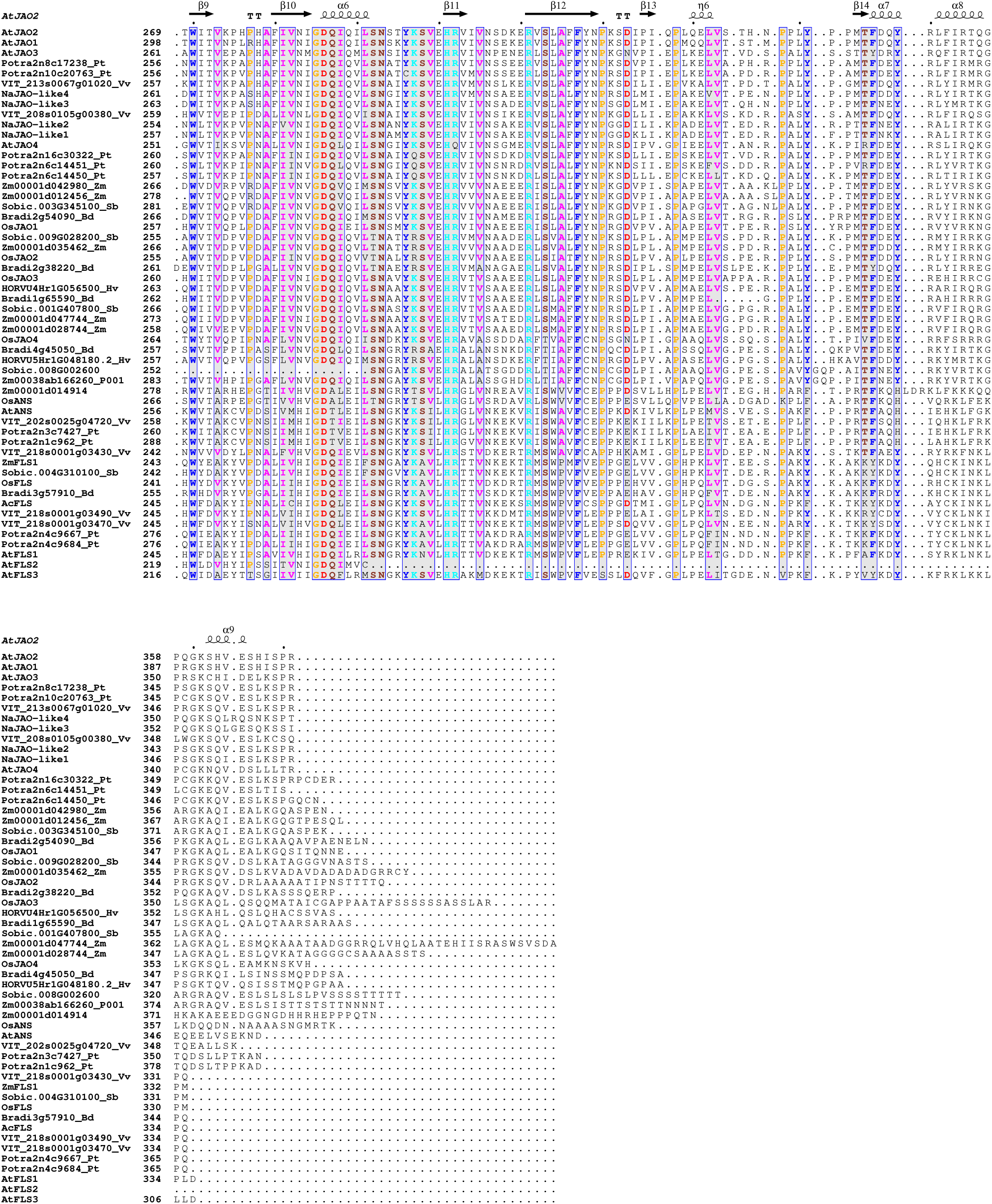
Multiple sequence alignment of selected AtJAO2-related protein sequences from representative dicot and monocot species. Positions that are identical are highlighted by white letters and colored background; positions with >70% similarity are highlighted with a grey background. Letter colors indicate chemical family of residues: red: negatively charged; turquoise: positively charged; pink: hydrophobic; blue: aromatic; brown: non-charged lateral chain; orange: other. Alignment was performed with MUSCLE (Edgar, 2004) and displayed using the ESPript server (Robert and Gouet, 2014). Protein data entry for AtJAO2 is 6LSV. Supports Figure 1C. **Edgar, R.C.** (2004). MUSCLE: multiple sequence alignment with high accuracy and high throughput. Nucleic Acids Res **32**: 1792–1797. **Robert, X. and Gouet, P.** (2014). Deciphering key features in protein structures with the new ENDscript server. Nucleic Acids Res. **42**: W320–W324.

**Supplementary Figure S2.**
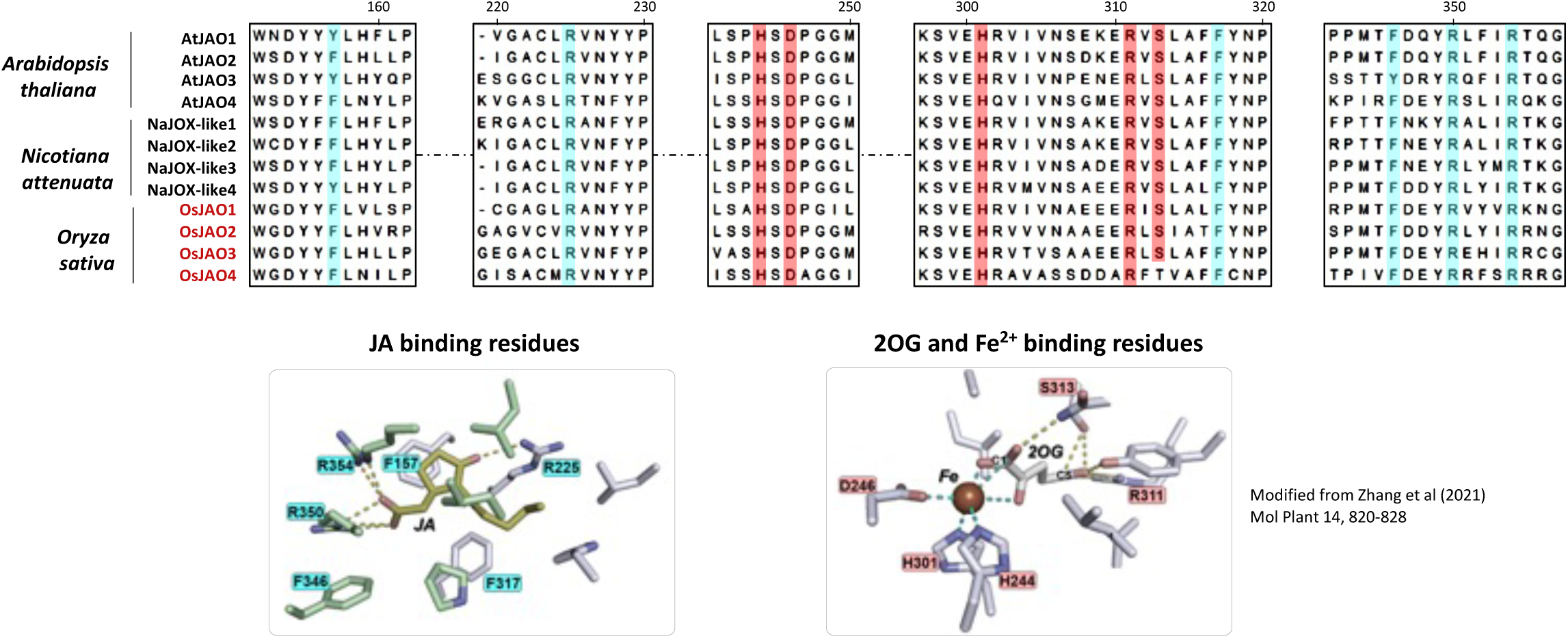
Alignment of predicted OsJAO protein sequences with partial sequences of functionally characterized JAO proteins from Arabidopsis and *Nicotiana attenuata*. Conserved amino acid residues identified by Zhang et al (2021) as essential for JA binding are highlighted in blue background; amino acid residues required for binding of co-substrate 2-oxoglutarate and iron are shown in red. The lower panels show the positions of conserved residues relative to JA substrate (left panel) and 2-oxoglutarate co-substrate (right panel) in the structure of the active site proposed by Zhang et al (2021).

**Supplementary Figure S3.**
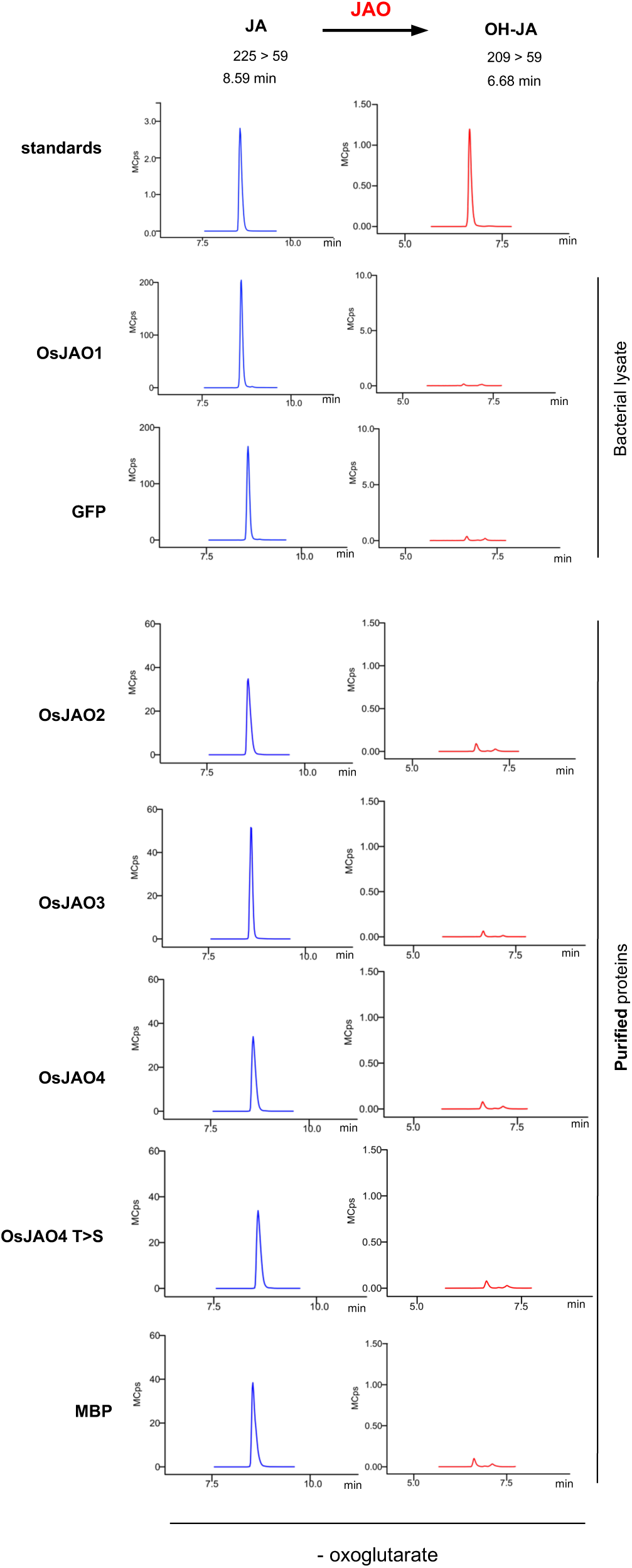
Negative controls for incubation reactions of recombinant OsJAO from bacterial lysate (OsJAO1 and GFP) or affinity-purified proteins (OsJAO2, 3 and 4) shown in Figure 2a. Reaction mixtures omitted the co-substrate 2-oxoglutarate. Reaction mixtures were analyzed by LC-MS/MS.

**Supplementary Figure S4.**
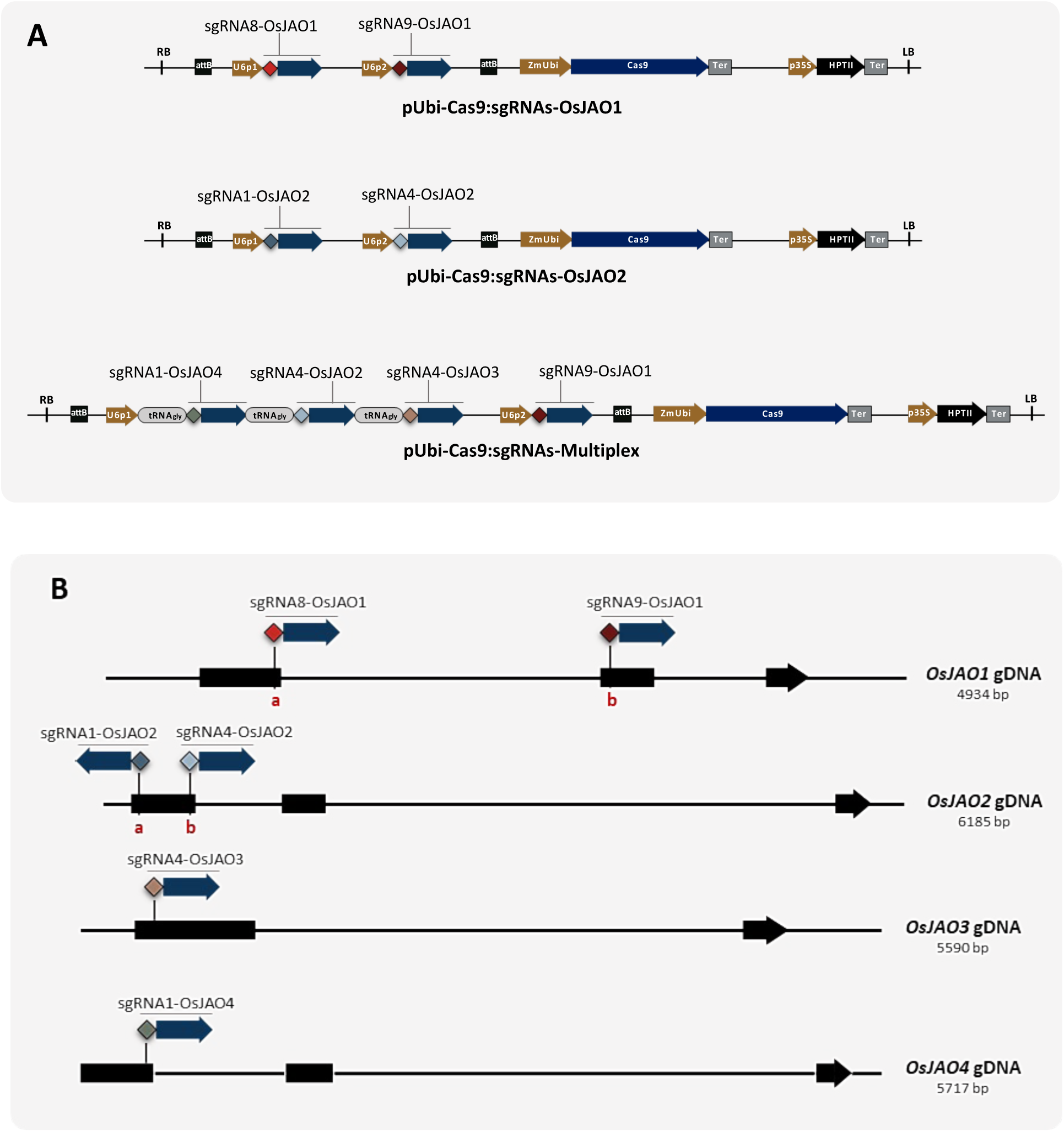
CRISPR-Cas9 plasmid constructs used for generating *Osjao* mutant lines and positions of targeted sequences for each guide RNA (sgRNAs) on *OsJAO* genes. (a) Graphical charts of T-DNA regions of the three constructs used to express the mutagenesis CRISPR-Cas9 machinery (Cas9 and gRNA) to produce single *jao1* and *jao2*, and multiple *jao* rice mutant lines. On pUbi-Cas9:sgRNAs-OsJAO1 and pUbi-Cas9:sgRNAs-OsJAO2 constructs, the two sgRNAs selected for the gene were placed under the control of distinct Ubiquitin-6 promoters (U6p1 et U6p2 respectively). On the pUbi-Cas9:sgRNAs-Multiplex construct, the sgRNA-tRNA^gly^ multiplex module was placed under the control of U6p1 for OsJAO2, 3 and 4 and U6p2 for OsJAO1. (b) Graphical chart of *OsJAO1*, *OsJAO2*, *OsJAO3*, *OsJAO4* genes, showing the positions of target sites of each sgRNA on the respective genes. Genes are represented by filled rectangles (exons) and solid lines (non-coding sequences). Oriented sgRNAs are depicted by filled arrows and diamonds.

**Supplementary Figure S5.**
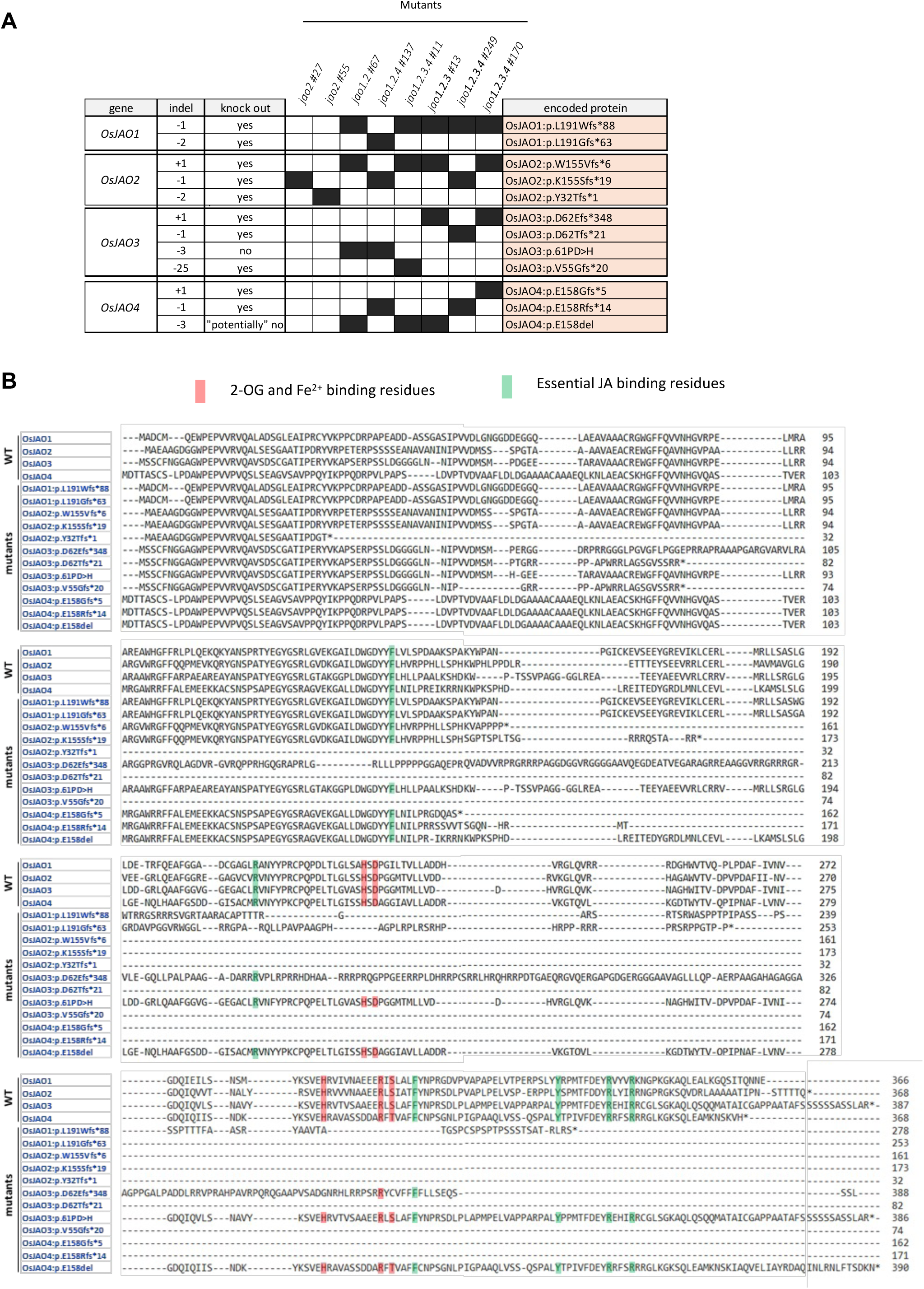
Representation of mutations (indels) identified in *OsJAO* genes in different rice mutant lines in T1 generation, and truncated proteins encoded by these altered genes. (a) Summary of the nature and combinations of indels in the series of single and multiple *Osjao* mutant lines obtained. The column ‘indel’ lists the indels detected and the filled rectangles indicate their occurrence in different mutant lines. In black: stabilized indel (homozygote). The right column describes the impact of the mutation on the structure of a putative encoded protein, using the nomenclature of Escande and Rouleau (2015). fs: frameshift. del: deletion. The number after the asterisk indicates the number of amino acid residues from the frameshift to the next stop codon (b) Alignment of wild-type OsJAO protein sequences with those of their mutant variants potentially expressed in obtained *Osjao* mutant lines. Predicted protein sequences derived from mutated gene sequences were aligned using ‘Muscle’ software to visualize impact of mutations on protein primary structure. Residues required for JAO activity (2-OG/Fe2+ binding and essential JA binding residues) are depicted with pink and green background respectively and can be checked for presence/absence in mutant proteins.

**Supplementary Figure S6.**
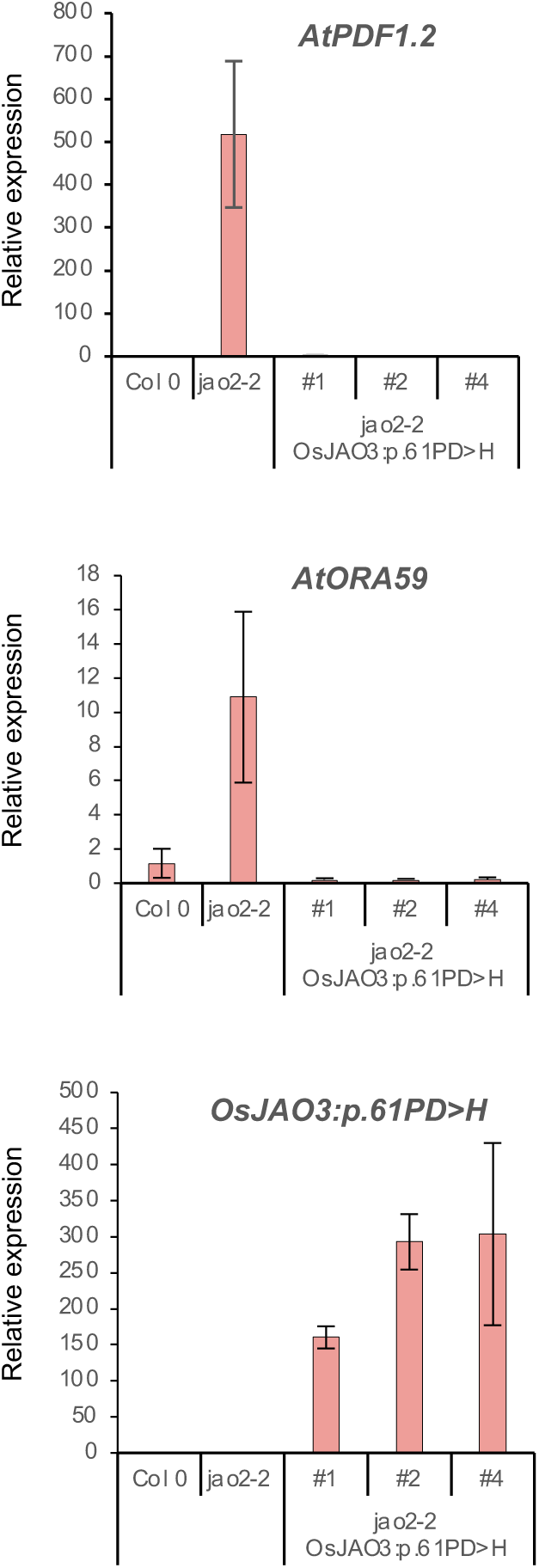
*In planta* assay of OsJAO3:p.61PD>H functionality. The cDNA encoding OsJAO3:p.61PD>H was cloned from the mutant rice line *jao1.2 #67* and ectopically expressed in Arabidopsis *jao2-2* line under p35S promoter. RNA from three independent T2 transformants along with untransformed WT (Col-0) and *jao2-2* was analyzed for *PDF1.2* marker gene expression. Expression was normalized with signal from *EXP* and *TIP41* housekeeping genes. Histograms show mean ± SEM from three biological replicates.

**Supplementary Figure S7.**
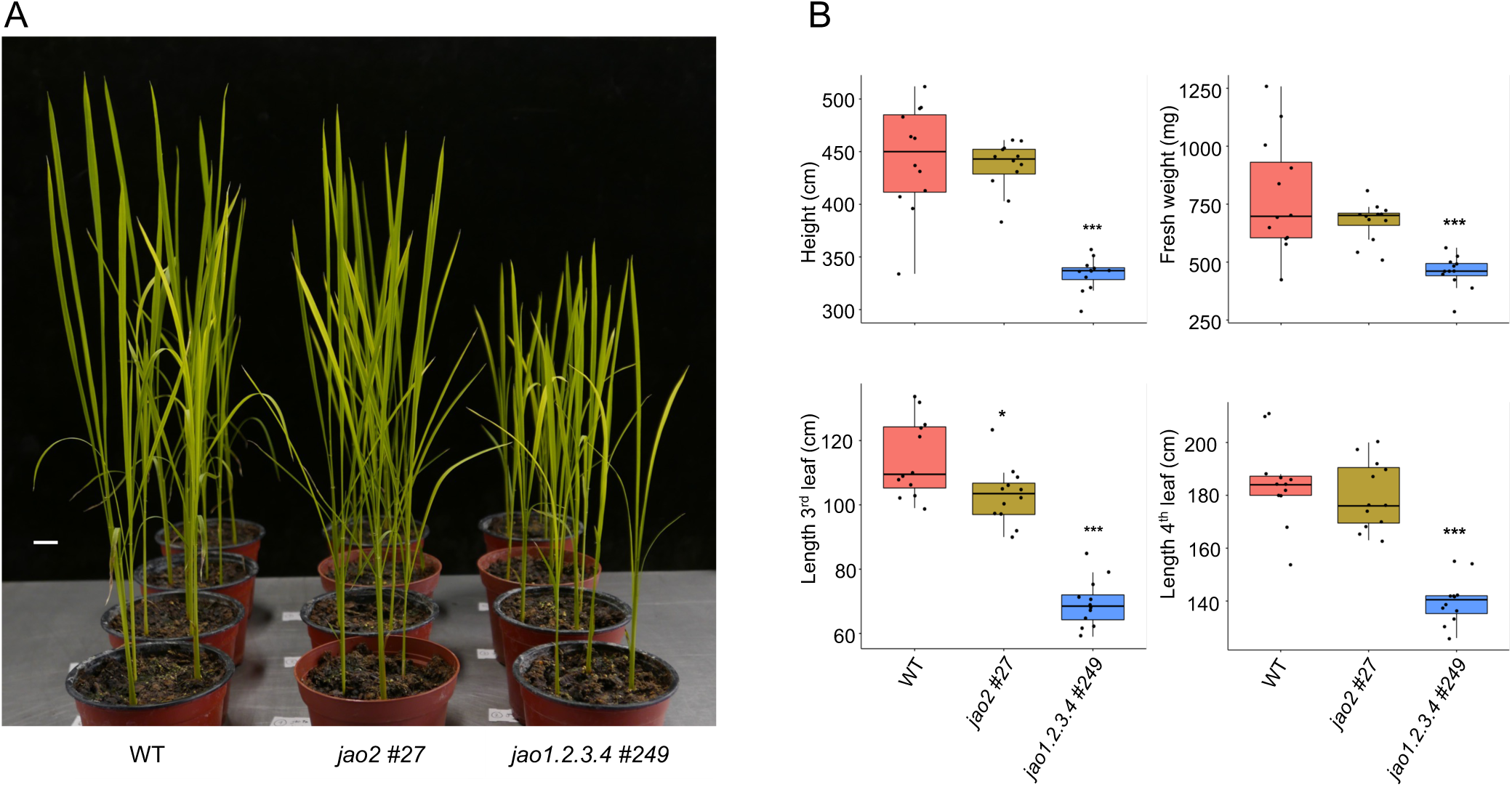
Analysis of shoot growth parameters of rice mutant lines impaired in OsJAO expression in T2 generation. (a) Picture of representative plants after four weeks of development. Scale bar for front row: 2 cm. (b) Shoot growth parameters of populations of WT, single *jao2* #27 and quadruple *jao1.2.3.4 #249* rice mutants (n= 12). Indicated genotypes were assessed for shoot height, shoot fresh weight and 3^rd^ leaf length. Asterisks indicate statistical difference with WT (t-test, \*\**P*<0.01; \*\*\**P*<0.001).

**Supplementary Figure S8.**
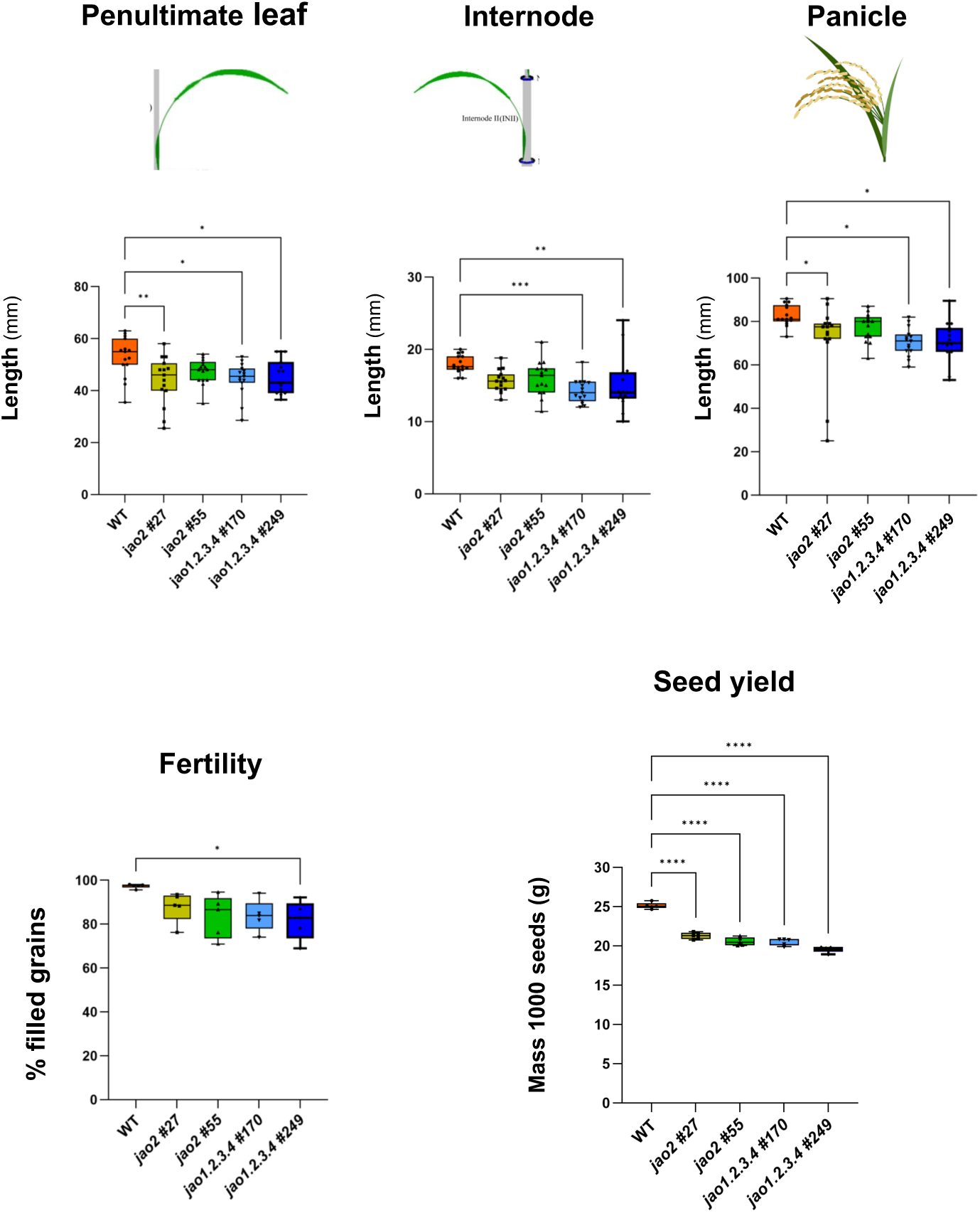
Analysis of shoot growth parameters of rice mutant lines impaired in *OsJAO* expression at T3 generation after 3 months grown in a greenhouse under a 16h-day/8h-night photoperiod, at 28°C/24°C with a 75% relative humidity. Penultimate leaf, internode and panicle lengths were measured at the mature stage on the 3 highest tillers from five T3 plants for each homozygous osjao line and wild type (WT) plants. Fertility rate was measured on the same tillers as the ratio between the number of fertile spikelets and the total number of spikelets. Seed yield was measured on the same tillers as the mass of 50 seeds of 15 panicles per genotype. Asterisks indicate statistical difference with WT (t-test, \*\**P*<0.01; \*\*\**P*<0.001; \*\*\*\**P*<0.0001).

**Supplementary Figure S9A.**
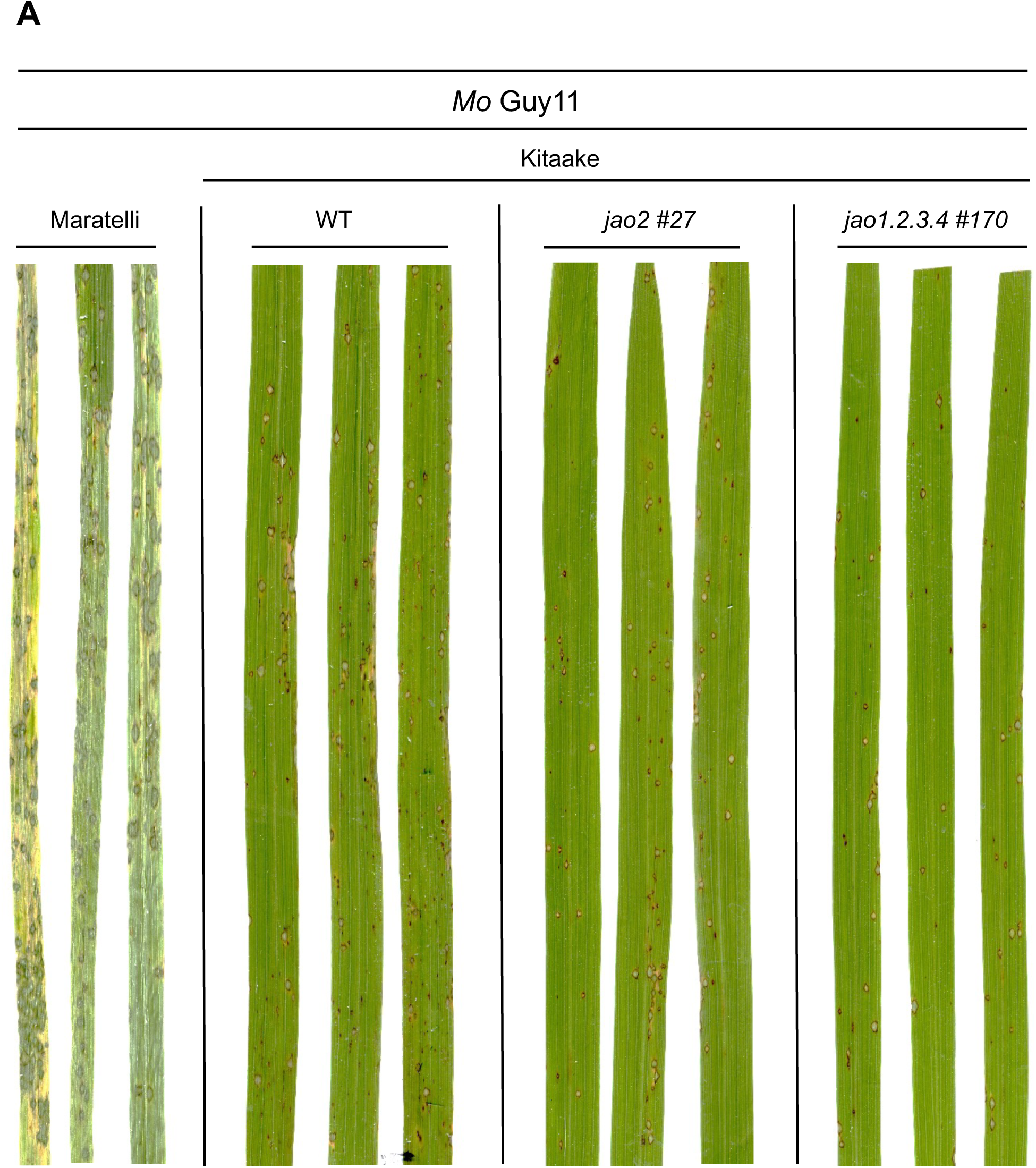
Representative images of leaves from Maratelli WT, Kitaake WT and Kitaake *jao* mutant infection experiment with *Mo* strain Guy11. For details see legend of Figure 7.

**Supplementary Figure S9B.**
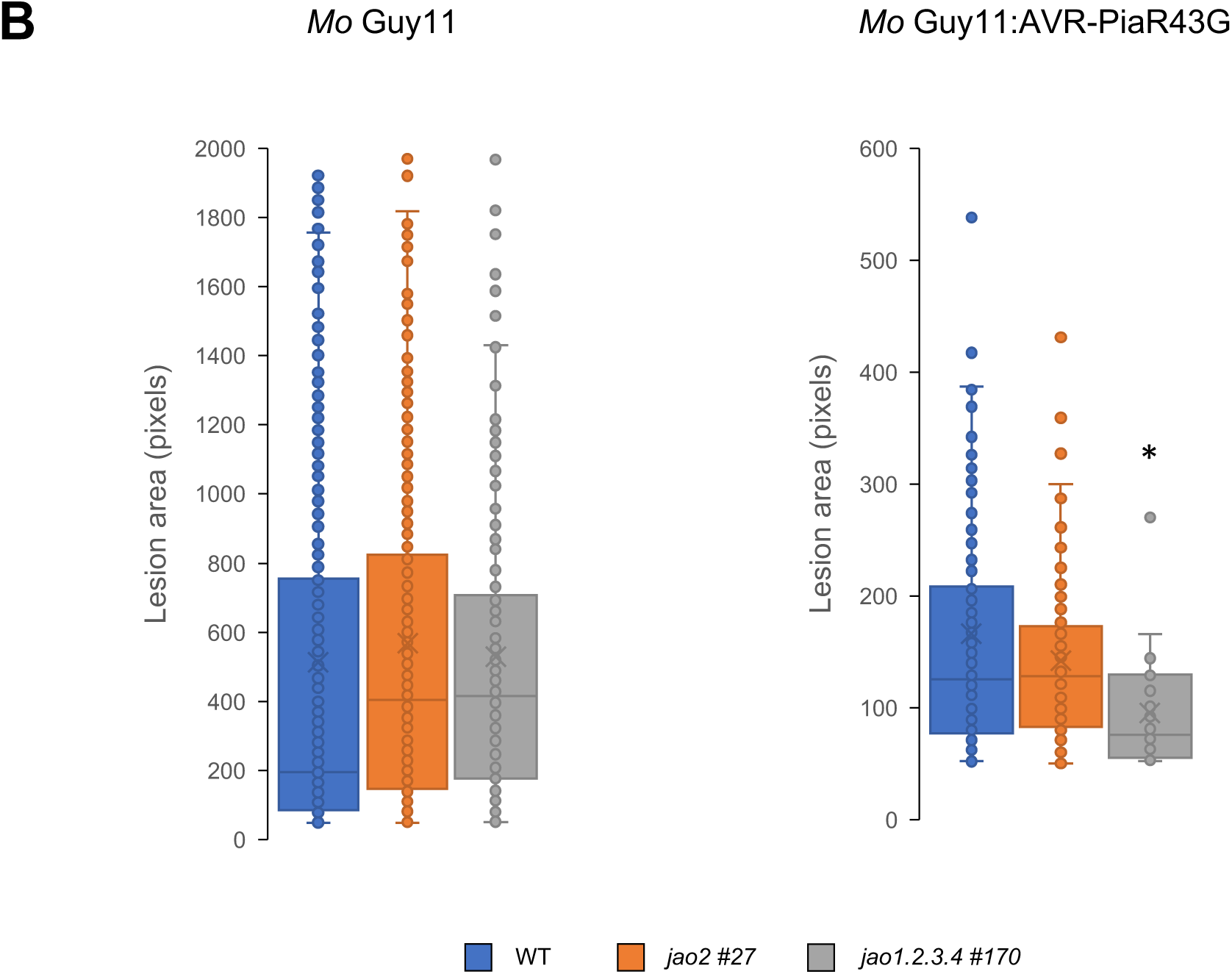
Individual lesion area on leaves of plants inoculated with strains *Mo* Guy11 (left panel) or *Mo* Guy11:AVR-PiaR43G (right panel) 7 days after inoculation. The boxes represent the second quartile, median, and third quartile. Individual data points are presented as circles and mean values as crosses. The mean values of the *jao1.2.3.4 #137* mutant were according to a t-test significantly different from those of the Kitaake WT with p<0,005 and are marked with *.

**Supplementary Figure S9C.**
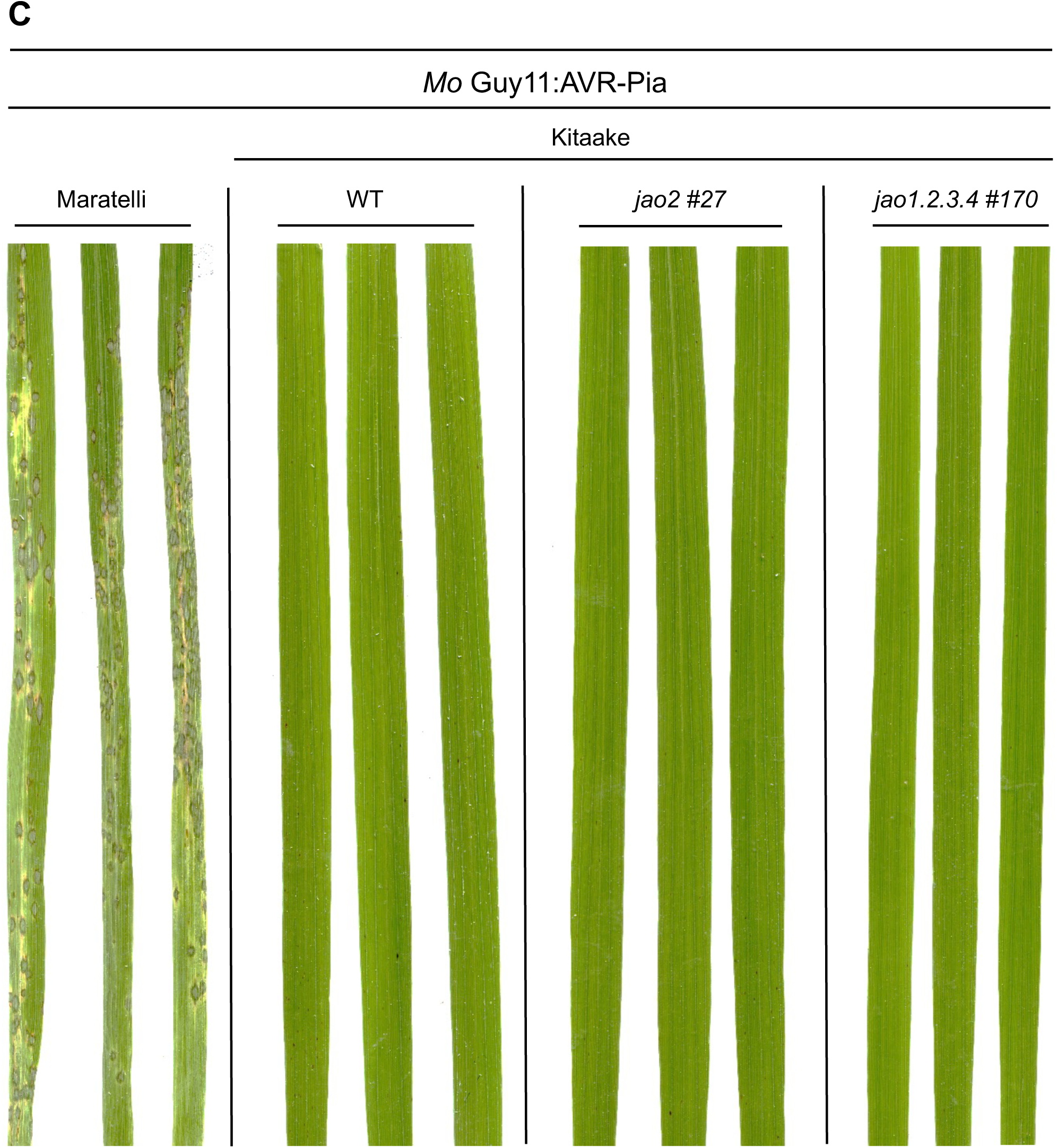
Representative images of leaves from Maratelli WT, Kitaake WT and Kitaake *jao* mutant infection experiment with *Mo* strain Guy11:AVR-Pia. For details see legend of Figure 7.

**Supplementary Figure S9D.**
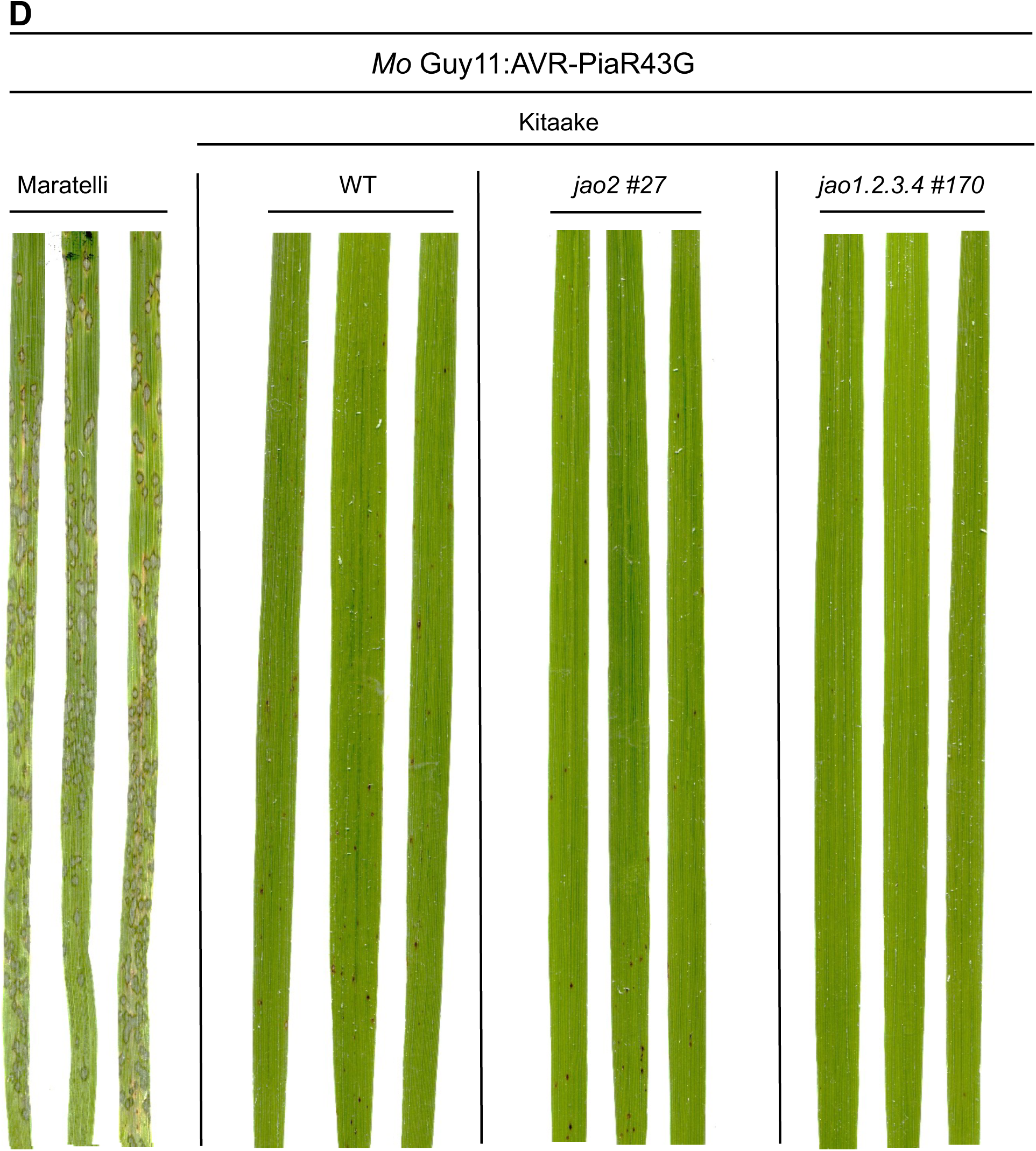
Representative images of leaves from Maratelli WT, Kitaake WT and Kitaake *jao* mutant infection experiment with *Mo* strain Guy11:AVR-PiaR43G. For details see legend of Figure 7.

**Supplementary Figure S9E.**
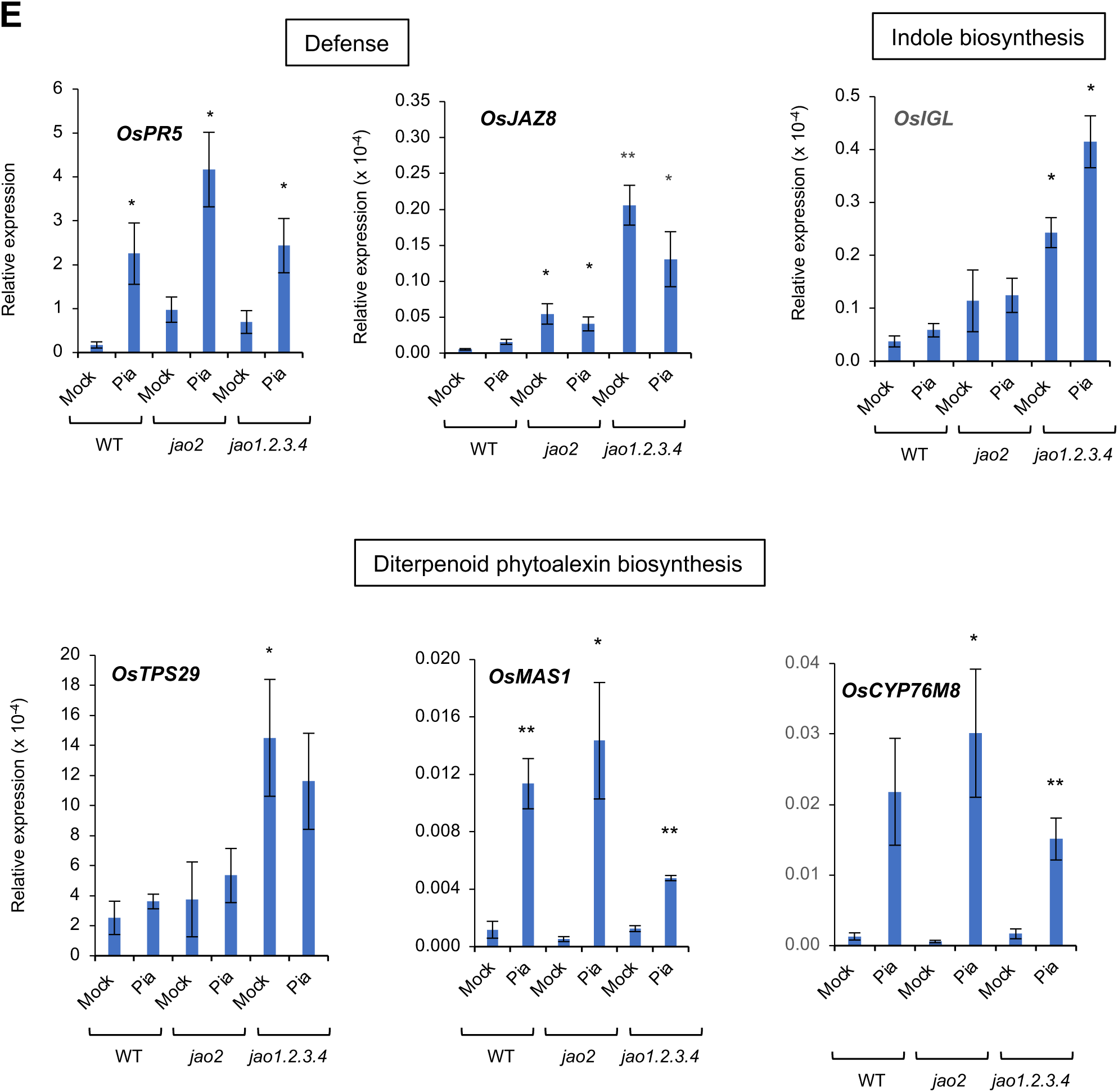
Expression of selected defense-related genes in *jao* mutant plants at 48 h after mock- or Mo Guy11:AVR-PiaR43G- (Pia) inoculation. *OsPR5*: Pathogenesis-Related 5; *OsJAZ8*: Jasmonate ZIM domain 8; *OsIGL*: indole glycerolphosphate lyase; *OsTPS29*: Terpene synthase 29; *OsMAS1*: momilactone synthase 1; *OsCYP76M8*: cytochrome P450 76M8. Expression was normalized with signal from *UBQ5* and *UBQ10* housekeeping genes. Histograms represent means of 3 biological replicates with SEM. Asterisks represent statistical significance for each condition relative to WT Mock (t-test, \**P*<0.05; \*\**P*<0.01).

**Supplementary Table S1.:** Metabolic features detected in extracts of 9-d old rice seedlings. Features are named by their m/z value and tentatively annotated by database interrogation as described in Methods section. Signal intensity is given for each biological replicate in indicated genotypes. RT: retention time.

**Supplementary Table S2.** Differential compounds (DC) between each of 8 JAO-deficient mutant lines and WT Kitaake. Leaves of 9-d old plants were analyzed (n=5). Compounds are named (ID) by m/z value and tentatively annotated after interrogation of indicated databases as described in Methods section. Color code refers to fold change (FC) relative to WT after pairwise comparisons with p value > 0.1.

**Supplementary Table S3.**
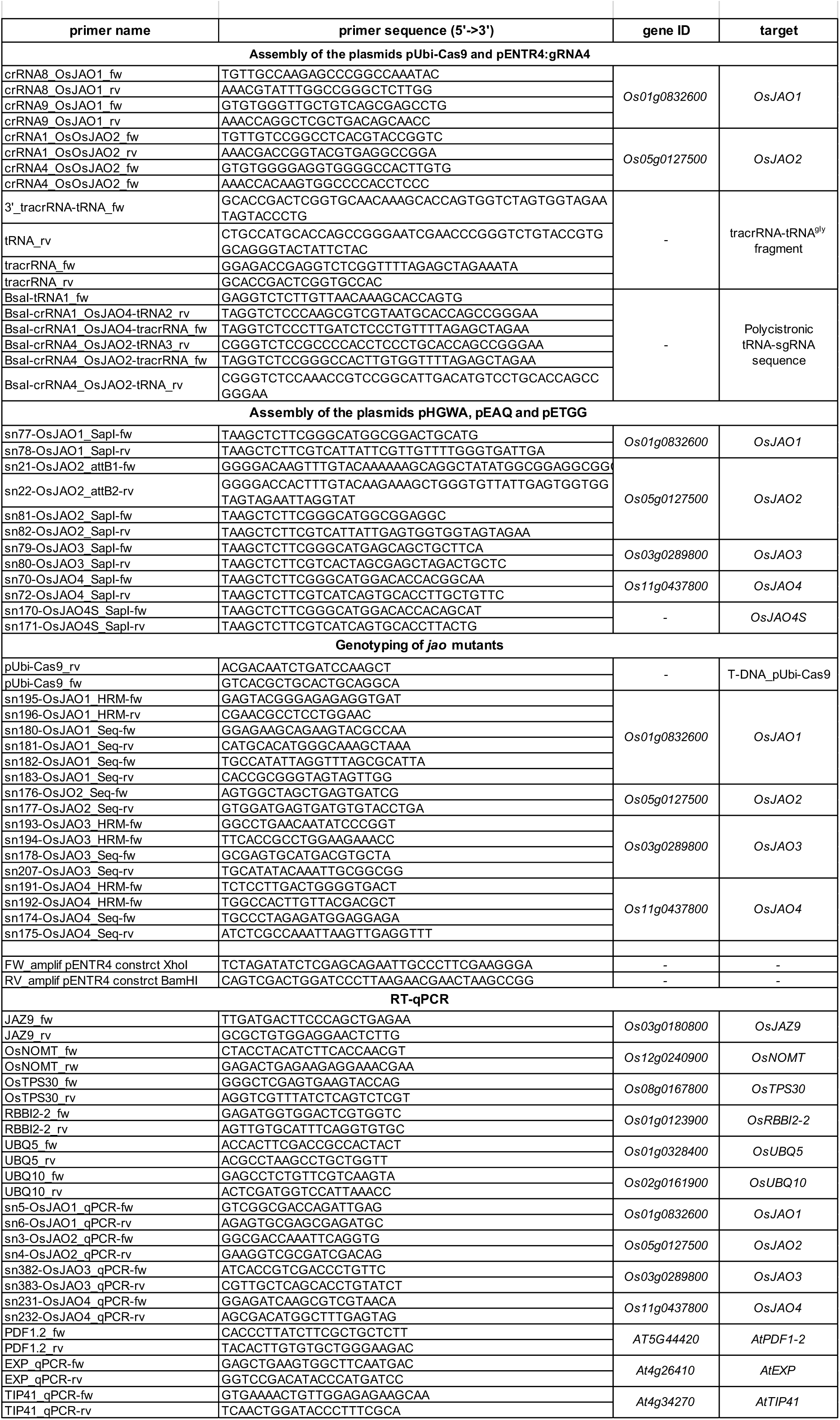
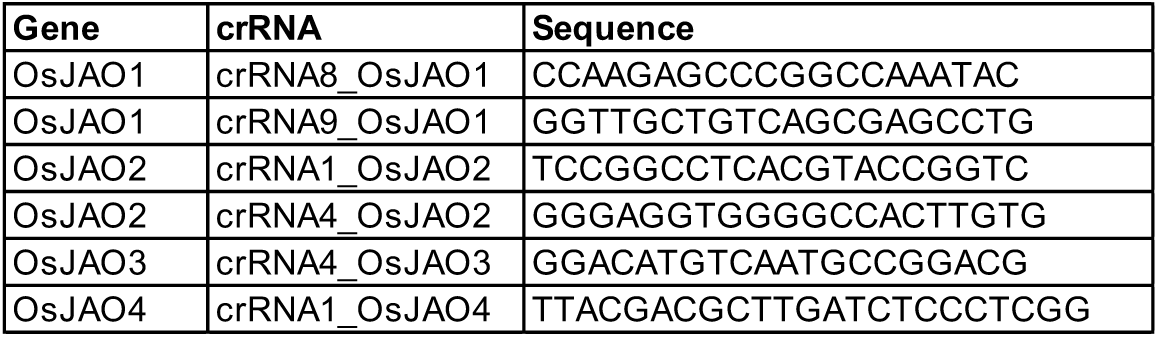
Primers used in the study.

## Notes

### Competing Interest Statement

The authors have declared no competing interest.

